# Is Over-parameterization a Problem for Profile Mixture Models?

**DOI:** 10.1101/2022.02.18.481053

**Authors:** Hector Baños, Edward Susko, Andrew J. Roger

## Abstract

Biochemical constraints on the admissible amino acids at specific sites in proteins leads to heterogeneity of the amino acid substitution process over sites in alignments. It is well known that phylogenetic models of protein sequence evolution that do not account for site heterogeneity are prone to long-branch attraction (LBA) artifacts. Profile mixture models were developed to model heterogeneity of preferred amino acids at sites via a finite distribution of site classes each with a distinct set of equilibrium amino acid frequencies. However, it is unknown whether the large number of parameters in such models associated with the many amino acid frequency classes can adversely affect tree topology estimates because of over-parameterization. Here we demonstrate theoretically that for long sequences, over-parameterization does not create problems for estimation with profile mixture models. Under mild conditions, tree, amino acid frequencies and other model parameters converge to true values as sequence length increases, even when there are large numbers of components in the frequency profile distributions. Because large sample theory does not necessarily imply good behavior for shorter alignments we explore performance of these models with short alignments simulated with tree topologies that are prone to LBA artifacts. We find that over-parameterization is not a problem for complex profile mixture models even when there are many amino acid frequency classes. In fact, simple models with few site classes behave poorly. Interestingly, we also found that misspecification of the amino acid frequency classes does not lead to increased LBA artifacts as long as the estimated cumulative distribution function of the amino acid frequencies at sites adequately approximates the true one. In contrast, misspecification of the amino acid exchangeability rates can severely negatively affect parameter estimation. Finally, we explore the effects of including in the profile mixture model an additional ‘F-class’ representing the overall frequencies of amino acids in the data set. Surprisingly, the F-class does not help parameter estimation significantly, and can decrease the probability of correct tree estimation, depending on the scenario, even though it tends to improve likelihood scores.

---

Phylogenetic methods have been used to resolve many deep phylogenetic problems in the tree of life (Brown et al. (2013); Daubin (2002); Pisani et al. (2015); Raymann et al. (2015); Wickett et al. (2014)). To increase phylogenetic signal and decrease topological estimation variability, many orthologous genes are usually analyzed. These analyses involve: (1) the alignments being concatenated into a ‘super-matrix’ from which trees are estimated; or (2) estimation of individual gene/protein trees first which are estimated and then combined using super-tree or ‘species tree’ methods; or (3) simultaneous co-estimation of gene and species trees (Bryant and Hahn, 2020). Regardless of the method, as more genes or proteins are considered, systematic biases can become predominant factors affecting estimation (Philippe et al. (2011)), underscoring the importance of adequately modeling the nucleotide or amino acid substitution process.

The substitution process of amino acid sequences is usually modeled as a site-independent Markov process on a tree. The most common approach assumes constant stationary (i.e. equilibrium) frequencies of the amino acids and a constant matrix of exchangeabilities throughout the tree. The amino acid frequencies are usually estimated from the observed frequencies in the entire alignment. The matrix of exchangeabilities is fixed *a priori*, chosen from a set of empirically defined matrices, see for example (Jones et al., 1992; Whelan and Goldman, 2001; Le and Gascuel, 2008). Also, it is customary to account for different sites evolving at different rates (see for example (Yang, 1994)). These models with the same substitution process at each site except for distinct rates at sites, are known as *site-homogeneous models* (although these are not strictly homogeneous because of site-rate variation).

However, there are different ranges of amino acids admissible at sites in proteins because of functional or structural restrictions (Franzosa and Xia (2009); Goldstein (2008); Pál et al. (2006)). These ranges can vary widely, from a few or just one, to essentially all possible amino acids at a site (Halpern and Bruno (1998); Lartillot et al. (2007); Lartillot and Philippe (2004); Wang et al. (2008)). Consequently, site-homogeneous models, which overlook this across-site amino acid frequency heterogeneity, are less biologically plausible and are prone to long-branch attraction (LBA) artifacts (Feuda et al. (2017); Lartillot et al. (2007); Simion et al. (2017); Wang et al. (2008); Williams et al. (2013)). LBA is a pervasive systematic bias in tree estimation whereby distantly-related groups with long branches are artefactually grouped together (Felsenstein (1978); Philippe and Laurent (1998); Bergsten (2005)). Given that LBA is by far the most widely noted empirical pitfall of analyses that ignore rate or site-frequency heterogeneity (Brinkmann et al., 2005; Bergsten, 2005; Philippe et al., 2005; Qu et al., 2017), we focus on it herein.

Partition (Lanfear et al. (2016); Pupko et al. (2002); Yang (1996)) and mixture models (Lartillot and Philippe (2004); Le and Gascuel (2008); Schrempf et al. (2020); Si Quang et al. (2008); Wang et al. (2008)) have been used to model heterogeneity of the amino acid substitution process across sites. Both partition and mixture models allow some subset of parameters to vary over the alignment; in this study, the stationary frequencies of amino acids at sites is the focus of attention. For partition models, these are treated as constant within partitions and estimated separately for individual partitions with the partitions in these alignments frequently corresponding to different proteins. For mixture models, the parameter subsets vary over sites, and rather than estimating separate parameters for separate sites, they are modeled as being independent and identically distributed from a mixing distribution that is estimated.

The CAT model (Lartillot and Philippe (2004)), a popular Bayesian mixture model, was shown to be less prone to LBA artifacts and to fit data better than site-homogeneous models (Lartillot et al. (2007)).In this model, all amino acid profiles (also known as frequency vectors, or frequency classes) are assumed to be independent and identically distributed from a discrete distribution with a finite number of profiles. The prior for the profiles in this mixing distribution is given by a Dirichlet process model which, because the number of profiles can get arbitrarily large, effectively allows non-parametric estimation of the mixing distribution. Markov chain Monte Carlo (MCMC) techniques are used to jointly infer the number of frequency vectors, the frequency vectors themselves, the affiliations of each site to a given class, the rates at each site, the branch lengths, and the tree topology. Unfortunately, in its current implementation (Lartillot et al. (2013)), convergence may not be achieved in practice for data sets with large numbers of sites and taxa. In the maximum likelihood framework, models accounting for site heterogeneity include mixtures of substitution rate matrices predefined for sites coming from different secondary structural elements and surface accessibility classes (Goldman et al. (1998); Le and Gascuel (2008, 2010)), for different site rates (Le et al. (2012)), a mixture of distinct sets of branch lengths (Crotty et al. (2019)), and mixtures of amino acid site frequency profiles (Schrempf et al. (2020); Si Quang et al. (2008); Wang et al. (2008, 2014)). The latter, together with the CAT model, are known as *profile mixture models*, and have become widely used for analyses of deep phylogenetic problems.

For a frequency mixture, we refer to the set of stationary amino acid frequency profiles (frequency vectors for short) with their corresponding weights as a *mixing distribution*, and we refer to the indices of frequency vectors in the mixing distribution as the *classes*. Fixed frequency vectors are often used to reduce the complexity and computational cost of estimation with profile mixture models. These come from mixing distributions previously estimated from data bases of alignments (Schrempf et al., 2020; Si Quang et al., 2008; Wang et al., 2008). Similar to the empirical estimates of rate matrices, such mixing distributions are estimated from large data sets such as those described in Dufayard et al. (2005) and Sander and Schneider (1994). The techniques used to obtain empirical estimates of these mixing distributions vary. For example, in Si Quang et al. (2008) the authors introduced six mixing distributions having 10, 20, 30, 40, 50, and 60 classes that were estimated from large data sets by maximum likelihood estimation. These are known as the C10-C60 (or generically CXX) mixing distributions. Schrempf and colleagues (Schrempf et al. (2020)) used K-means and the CAT model to estimate empirical mixing distributions ranging from 4, 8, 16, up to 4096 classes. These are known as the universal distribution mixture, or UDM mixing distributions. All these models have pre-estimated frequency vector weights, but these are often re-estimated with the data in hand. In contrast, their frequency vectors are always fixed.

Profile mixture models are less susceptible to LBA than site-homogeneous models (Phillips et al. (2004); Lartillot et al. (2007); Wang et al. (2008); Philippe et al. (2009)). An ongoing concern articulated in several studies (e.g. Anderson and Lindgren (2021); Li et al. (2021)) is to what extent profile mixture models with a large number of extra classes that are not represented in the data result in estimates of quantities of interest such as topologies or edge-lengths that are excessively variable (i.e. how over-parameterization affects estimation). If excess variability is to be expected, it would imply that such profile mixture models should be avoided in favor of simpler models.

Over-parameterization has been an active concern in phylogenetic studies, see for example (Lemmon and Moriarty, 2004; Huelsenbeck and Rannala, 2004; Sullivan and Joyce, 2005; Zhou et al., 2007; Groussin et al., 2013; A Shepherd and Klaere, 2018; Guimarães-Fabreti and Höhna, 2022). Part of our investigation includes the formulation that profile mixture models have desirable properties when inferring parameters. Particularly, as carefully described in the next section, we argue that by combining the results in Kiefer and Wolfowitz (1956); Lindsay (1983), tree and mixing parameters are statistically consistent even with a large number of profiles; that is, parameter estimates are expected to get arbitrarily close to true parameters as the number of sites increases.

Although the foregoing consistency property is desirable in any modeling context, there is no guarantee that such models will have good small sample properties. Thus there is a need to explore the performance of profile mixture models through simulations of short alignments both with and without model misspecification.

In this work we consider simulations from empirically-derived mixtures resulting in alignments with lengths ranging from 150 to 1000 sites. Of particular interest was accuracy and variability of topology estimation as well as the accuracy of estimation of the mixing distribution. Accuracy of mixing distribution estimation was measured by a distance between the estimated and true cumulative distribution functions (CDFs) of the frequency vectors.These CDFs, properly described in the following section, are an alternative way to re-parameterize profile mixture models. Other works, for example (Luo et al., 2010; Abadi et al., 2019; Zou et al., 2019; Zaharias et al., 2022), have analyzed the performance of complex models with short alignments. Unlike these studies, here we focus specifically on profile mixture models and the potential effects of over-parameterization.

We also explore the effects of the “F-class,” a class that is defined from the empirical frequencies of amino acids from the overall alignment, that is often included as an additional class in models to account for remaining sites in the data that are not well modeled by the fixed empirically-derived frequency vectors. Finally, we explore the performance of profile mixture models on three empirical data sets. The goal of these is to investigate the performance of different models and the F-class for long alignments in the presence of model misspecification.

## Materials and Methods

### Over-parameterization

Over-parameterization generally refers to model formulations involving a large number of parameters that result in poor statistical performance. The term is not well defined for several reasons. First, the aspect of statistical performance of concern depends on the context. In machine learning or regression modeling, where the goal is often prediction, models that are too parameter rich often predict the observed data well but are very poor at extrapolating to new data. When the parameters of the model are of primary interest, a model might be considered over-parameterized because a large number of parameters results in high parameter variance. For instance, regression models with many predictor variables often give extremely variable regression coefficient estimators. More subtly, a model might be considered over-parameterized when there is a poor statistical estimation of some specific quantity of interest separate from a large number of parameters themselves. It is these last two cases of over-parameterization that are of primary interest in phylogenetics. An example of the latter would be a parameter-rich phylogenetic model that gives poor topological estimation.

Another reason that over-parameterization is not well defined is that poor statistical performance exists on a continuum. The most unambiguous examples of over-parameterization are when estimators become statistically inconsistent for models with too many parameters. For instance, topology estimation can be statistically inconsistent for partition models with many partitions and a relatively small number of sites per partition (Tuffley and Steel, 1997; Roch et al., 2018; Wang et al., 2019; Susko and Roger, 2021). Statistical inconsistency is an extreme form of poor statistical performance. At the other extreme of poor statistical performance, one might consider a model to be over-parameterized if it gives more variable or more biased estimates than some other model with fewer parameters. Under these criteria, a phylogenetic model with a single set of free frequency parameters might be considered over-parameterized relative to a model where frequencies are all fixed to be equal. If the true model does indeed have equal frequencies, the model with free frequency parameters is not expected to perform as well. We adopt instead the perspective that over-parameterization requires not just that statistical performance to be poorer but that it be substantially poorer, with the term ‘substantially’ being admittedly vague. In what follows we find, however, that parameter-rich mixtures tend to perform better than simple models albeit, as expected, not better than the model where parameters are known.

The example above involving the estimation of a single set of frequencies underscores yet other reasons that over-parameterization is not well-defined. First, poor statistical performance requires a comparator method (for example, frequency estimation vs all equal frequencies). Second, statistical performance is dependent on a true model. When the true model has equal frequencies, models with frequency estimation are expected to perform more poorly than models with equal frequencies but if the true model does not have equal frequencies, the bias incurred by an equal-frequency fitted model might outweigh its smaller variance. In our examples, we include both cases where the true model is among the fitted models and where all of the fitted models are misspecified.

### Mixture models and over-parameterization with large samples

In this section, we define and elaborate on some theoretical properties of profile mixture models. Re-parameterizing the mixing distributions as the CDFs of the amino acid frequencies at a site provides insight into why over-parameterization is not a problem for profile mixture models. Also, we briefly discuss known identifiability results for such models.

Roughly speaking, profile mixture models are mixtures of time-reversible models, with a common exchangeability matrix *R*. The parameter space of a profile mixture model is usually defined by

i. A rooted metric tree *T* on *N* taxa.
ii. A symmetric 20 *×* 20 matrix of non-negative exchangeabilities *R*.
iii. A collection of *C* frequency vectors {***π***_*c*_}, and corresponding weights *{w*_*c*_*}* with *w*_*c*_ ⩾ 0 and 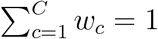. Together these components give the profile mixing distribution of the profile mixture model.
iv. A collection of *K* scalar rate parameters *{r*_*k*_*}*, with *r*_*k*_ ⩾ 0, and rate weight *d*_*k*_, with *d*_*k*_ *>* 0 and 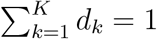.

The substitution process of a profile mixture model is as follows: a frequency vector is sampled from the mixing distribution, such that it is equal to ***π***_*c*_ with probability *w*_*c*_, and a rate parameter is sampled from the rate distribution, such that it is equal to *r*_*k*_ with probability *d*_*k*_. Evolution of a sequence at a site is then according to a continuous Markov substitution process over tree *T* with exchangeabilities *R*, root distribution ***π***_*c*_, and rate *r*_*k*_. For a given site pattern ***x***_*i*_, the contribution to the likelihood is a weighted average of partial site likelihoods conditional on each site-profile class and site-rate class:

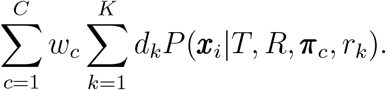

For a model with fixed amino-acid frequency classes, a natural way of parameterizing the mixture model is in terms of its weights, *w*_*c*_, *c* = 1, …, *C*. This leads to models of differing dimensions *C* that, as mentioned before, can get very large, raising concerns about over-parameterization.

An alternative way of parameterizing the mixture is in terms of its cumulative distribution function (CDF) of the amino acid frequencies at a site. Let **Π** = (Π_1_, …, Π_20_) denote the random frequency vector for a site. Then the CDF is

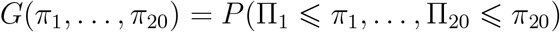

This allows one to express mixing distributions with differing components (*C* = 20 say or *C* = 60) as being in the same parameter space. Recall that the cumulative density function *F*_*X*_(*x*) of a random vector *X* = (*X*_1_, …, *X*_*q*_) evaluated at *x* = (*x*_1_, …, *x*_*q*_), is the joint probability that *X* takes values less or equal to *x*. That is *F*_*X*_(*x*) = *P* (*X*_1_ ⩽ *x*_1_, *X*_2_ ⩽ *x*_2_, …, *X*_*q*_ ⩽ *x*_*q*_) (see the Appendix for a review on CDFs). The CDF is uniquely determined by its weights and frequency vectors. Conversely, whenever it is a *finite mixing distribution*, meaning that it is describable in terms of a fixed set of weights and frequency vectors, the CDF can be used to determine what those weights and frequency vectors are.

One advantage of this parameterization is that distributions that are similar will have similar CDFs but might have very different *w*_*c*_ parameters or include some very different frequency vectors with small weights. We give an example in the Appendix.

However, now the frequency mixture parameter space, which is a space of distribution functions, is infinite-dimensional, and would appear to be an extreme case of over-parameterization.

Surprisingly, even for this infinite dimensional space of distribution functions, estimation of both the mixing distribution and structural parameters like the tree is frequently still consistent as was shown by Kiefer and Wolfowitz (1956) for a wide class of models under mild regularity conditions. Most of these conditions are expected to hold for the models considered here. A more detailed explanation of how the results in Kiefer and Wolfowitz (1956) apply to our context is given in the Appendix.

The implication of Kiefer and Wolfowitz (1956) is that class frequency mixture models are not over-parameterized, at least with large samples. The reason for this is that, here, the space of all distribution functions as a space of functions is relatively “small” in the mathematical sense of being a compact space (a closed and bounded space in our setting). An alternative way of seeing why this is the case is to note that the *w*_*c*_ are restricted to be non-negative and sum to one. By contrast in cases of classical over-parameterization, parameters usually are unrestricted (for example, a regression model where there are more predictors than observations).

Another surprising result of estimation within the mixing distribution setting is that even if parameter estimation is unrestricted and any mixing distribution is allowed, the maximum likelihood estimator will be a finite mixing distribution. This is an implication of the results of Lindsay (1983) as detailed in the Appendix. From this we can conclude that when the model is correctly specified and identifiable, one can effectively estimate the true parameters as the number of sites increases.

With respect to identifiability, Yourdkhani et al. (2021) report results for a large family of profile mixture models. These authors showed generic identifiability of the tree and mixing parameters for models with *C* · *K <* 72, where *C* is the number of classes and *K* the number of rates, trees with more than 8 taxa, and where no parameters are assumed to be fixed. As discussed in the Appendix, these results also apply to some of the models with fixed frequency vectors considered here. We conjecture that, generically, identifiability of the tree topology can be achieved for models with fixed frequency vectors with *C* classes, *K* rates and *m* taxa, for some *C* · *K >* 72, and all *m >* 8. This will be explored in future work.

### Simulation setting

In this section, we describe all the different parameters used to simulate alignments under profile mixture models. These are presented below in the following order: (1) the trees; (2) mixing distributions; (3) exchangeability matrices; (4) sequence lengths; and (5) rate parameters.

By choosing different combinations of parameters, we simulated a total of 396 scenarios. For each scenario, 100 simulations were performed using Alisim (Ly-Trong et al. (2021)). The choices of parameters were as follows.

### Trees

Nine different trees are considered for these simulations. All the trees have the ‘structure’ of tree *T* shown in Figure 1. The features that vary per tree are the length of a single edge *l*, where *l* ∈ *{*0.005, 0.02, 0.05*}* and the number of taxa at each polytomy *m*, where *m* ∈ *{*1, 2, 3*}*. We denote each of the trees by *T*_6*m*_(*l*).

**Fig. 1.**
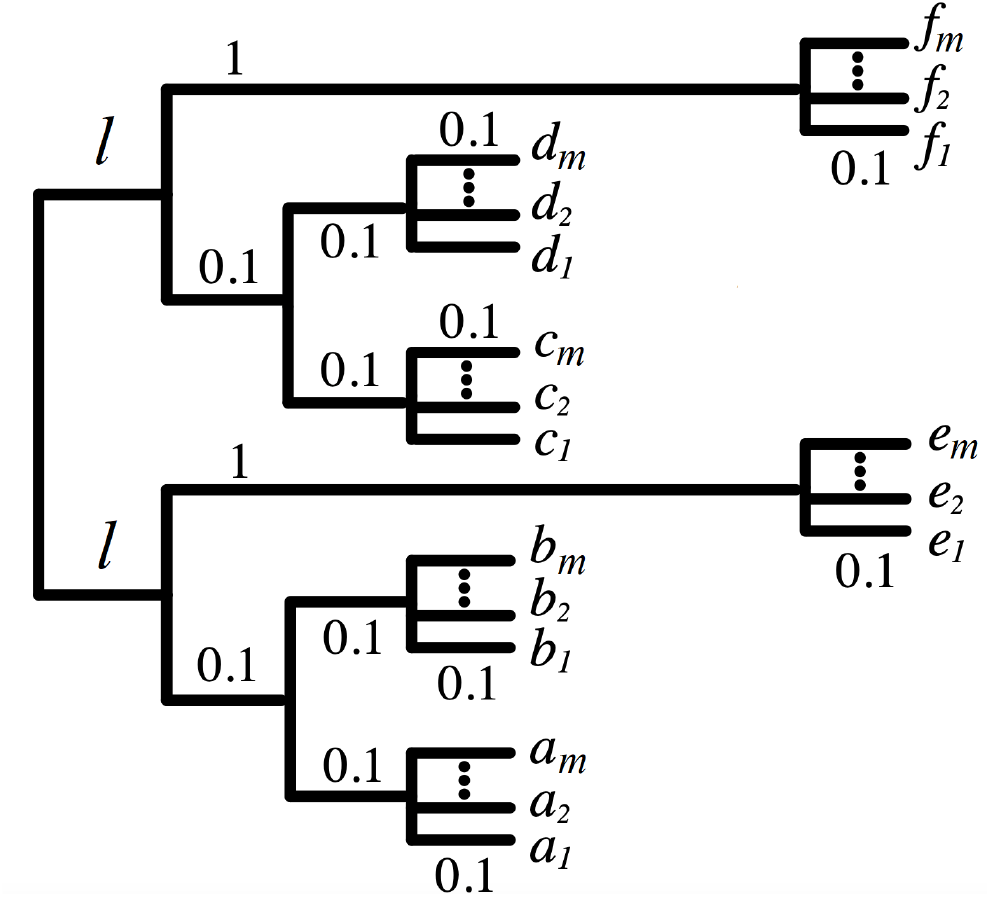
The main structure of the tree where all the simulations were conducted. Only the edge lengths *l* and the number of taxa at each polytomy *m* are variable. This tree has 6*m* taxa

The structure of *T* is chosen since it is often a tree susceptible to LBA artifacts. For fixed *l* and changing *m*, the simulations are, in effect, all from the same tree but with differing levels of taxonomic sampling from the 6 clades. By increasing the number of taxa, we obtain more information on the frequency vectors. By decreasing the edge length *l*, we make the tree more susceptible to LBA artifacts whereby the f-clade (i.e. clade including taxa *f*_1_, *f*_2_, …, *f*_*m*_) and the e-clade (i.e. clade with taxa *e*_1_, *e*_2_, …, *e*_*m*_) group together to the exclusion of the other clades. Also, structurally, these trees are meant to show how well a split can be inferred under conditions where an LBA bias is very likely to arise.

We also conducted additional simulations on three larger trees and the results are reported in the *Supplementary material*. For these latter analyses, we used two empirically-derived trees with 40 and 32 taxa, and a tree simulated under a birth-death process on 40 taxa (see the Supplementary Material for the details on these trees).

### Mixing distributions

For each tree, we simulated data using ten different mixing distributions. One of these is the model C60 as defined in Si Quang et al. (2008), which has 60 classes. Another seven are built on the frequency vectors of C60. Specifically, for *i* ∈ *{*10, 15, 20, 30, 40, 50, 60*}*, we defined the mixing distribution C60[*i*] by choosing *i* frequency vectors from C60 at random and assigning them non-zero weights sampled from a Dirichlet distribution with concentration parameters ***α*** = 1. We denote by C60 the set of mixing distributions C60[*i*], for all *i*, including C60. The two remaining mixing distributions used to simulate data are known as UDM-0256, and UDM-4096 (Schrempf et al. (2020)), that have 256 and 4096 classes respectively. Specifically, we used the non-transformed mixing distributions denoted as UDM-0256-None and UDM-4096-None in Schrempf et al. (2020).

In reality, every site in a protein has a unique physicochemical environment. Although this environment may drift over evolutionary time as a result of substitutions at other sites, conservation of site preferences for amino acids is often observed in alignments (Youssef et al., 2022). Therefore profile mixture models with fixed frequency classes approximate these commonly occurring patterns amongst sites. By simulating under C60, the most complex of the CXX mixing distributions, and the complex UDM mixing distributions, we try to emulate real data. The C60[*i*] distributions reflect the scenario wherein, for a small sample, not all relevant frequency vectors are represented, nor are distributed as in C60. We note that the UDM mixing distributions are distinct from the CXX distributions but because the data-sets used to estimate both CXX and UDM mixing distributions overlap, some similarities in their frequency profiles are expected.

### Exchangeability Matrices, Sequence Lengths, and Site Rate Variation

For all combinations of mixing distributions and trees, we simulated data using the exchangeability matrix from the LG model (Le and Gascuel (2008)); we refer to this as the LG matrix. We also used a “POISSON” exchangeability matrix, a matrix with equal exchangeabilities, but only for simulations involving the two UDM mixing distributions. We did not simulate alignments using a POISSON matrix for the ℭ60 mixing distributions since there are many simulations settings being considered already.

For all combinations of mixing distributions, trees, and matrices, we used sequence lengths of 300, 600, and 1000 amino acids. In practice, alignments of length 300 are more typical for single protein data sets. For the mixing distributions in ℭ60 we also simulated sequences of length 150 to explore their behavior with smaller proteins.

Lastly, for all simulations we used four rate parameters coming from a discretized Γ(4) distribution (Yang (1994)) with *α* = 0.5.

### Fitted models and Precision of parameter estimation

In this section, we describe how different choices of fixed frequency vectors and exchangeabilities were fitted to the simulations. We also introduce four criteria used to evaluate model fitness.

When fitting a model to data, estimated parameters were obtained by maximum likelihood. IQ-TREE 2 (Minh et al. (2020)) was used to get the maximum log-likelihoods and the estimators (MLEs) for all simulations. The parameters that were optimized by IQ-TREE 2 are the tree topology, edge lengths, and weights of the frequency vectors. All other parameters including the frequency vectors, the matrix of exchangeabilities, and the rate parameters, were supplied as fixed values to IQ-TREE 2.

The LG matrix was the only matrix used to simulate data under the mixing distributions in C60. To these simulated data sets, we fit the LG matrix, the F-class, and various frequency vectors. Specifically, the frequency vectors fitted included C60, C40, C30, C20, defined in Si Quang et al. (2008), LG (Le and Gascuel (2008)), LG4X (Le et al. (2012)), as well as the frequency vectors in ℭ60 used to simulate the data. In these analyses, we explored mainly two things: (1) possible model over-parameterization effects from fitting more general models to short alignments; and (2) the consequences of misspecification of the frequency vectors by using models that have different frequency vectors than those of the generating model.

For data generated under the UDM mixing distributions, we fitted the exchangeability matrices POISSON and LG, with and without the F-class, and frequency vectors of C60, C40, C30, C20, LG4X (models C50 and C10 were excluded to simplify the case study). We also fitted the POISSON model for data generated under POISSON exchangeabilities and the LG model for data generated under LG exchangeabilities. In these analyses we primarily explored the effect of: (1) misspecification of the frequency vectors; (2) misspecification of the matrix of exchangeabilities; and (3) use of the F-class in estimation. Recall that only the mixing weights, the tree topology, and edge-lengths are optimized, everything else is fixed before the maximization of the likelihood.

For one of the criteria used to compare overall model performance (described below), information on the second ‘best’ topology (the runner-up) is needed. Therefore, technically, all topologies must be optimized to be compared. To reduce the computational burden, we only considered a subspace of tree space. In preliminary results, we computed the likelihoods of all 105 6-taxon unrooted topologies for data generated under *T*_6_(0.005). We noted that there were two tiers in terms of log-likelihood values; these occurred regardless of the generating and fitting models. One tier consists of the log-likelihoods of the 35 topologies shown in Table S1 in the Supplementary Material, and the other consists of the remaining 70 trees. The first tier showed significantly larger log-likelihood values, see Figure S1 in the Supplementary Material. The 35 topologies in the first tier are all the topologies displaying the embedded quartet tree *AB*|*CD*. This shows, as expected, that there are no problems with estimating the relationships amongst the taxa in this quartet. The main difficulty was instead determining the correct placement of the long branches because of the LBA-related artifacts.

Therefore, for simplicity, in all cases we restrict the tree space considered to just these 35 tree topologies by substituting *X* ∈ *{A, B, C, D, E, F }* by the adequate *m*-taxon polytomy. We also consider in this tree space the ‘star tree’ topology obtained from *T* in Figure 1 by setting *l* = 0.

We now proceed to introduce the criteria used to compare the overall model performance.

### Integrated Squared Error

As mentioned earlier, profile mixture models can be parameterized as CDFs of the amino acid frequencies at a site. Therefore, a reasonable way to measure the precision of parameter estimation is by comparing how closely the observed CDF resembles the true one.

To compare CDFs we used the *Integrated Squared Error* (ISE), which is defined as follows

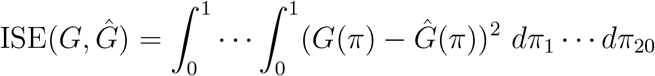

where *G* is the true CDF and *Ĝ*(*x*) is the CDF obtained from the estimated mixing distribution.

Because the ISE is a high-dimensional integral, it is not generally computational feasible. However, as we show in the Appendix, ISE can be calculated efficiently for the finite mixing distributions that are considered in phylogenetic applications.

For a given choice of parameters, we report the mean ISE for all 100 simulations. ISE satisfies the requirements of a distance. Thus ISE = 0 if and only if the two distributions are the same and approaches 0 if and only if the observed CDF converges upon the true. ISE cannot be used in practice since the true CDF is then unknown. Nonetheless, as detailed in the Results section, in this case, where the true CDF is known, it can effectively help us to assess model performance.

In order to assess the precision of parameter estimation in a more standard way, we also considered two additional criteria based on the tree topology estimate.

### Overall Accuracy and Proportional Mode

The following criterion, *overall accuracy* denoted OA, is a standard way to evaluate the precision of parameter estimation. Given a set of simulations, overall accuracy is defined as the proportion of times where the true tree was recovered (in other words, the proportion of times where the true tree topology was the one maximizing the likelihood).

The next criterion, the *proportion of settings where the true tree was the mode of the distribution of estimated trees*, (“proportional mode” for short) denoted PM, evaluates the precision of parameter estimation in a manner that can indicate whether there are biases in estimation. For a discrete distribution, the mode of the distribution is the outcome that comes up most frequently. For a given fitted model, we were interested in the proportion of parameter settings where the true topology was the mode of the distribution of the topology estimates for that model. This can be approximated by the proportion of simulation settings where, over repeated simulations, the estimated topology was equal to the true tree more frequently than any other particular tree. To account for some sampling variability in PM, in each scenario we tested if the proportion of times the true topology was chosen was significantly higher than another single topology via a binomial test of two proportions. That is, for each time the true tree was the most frequent in a setting (all 100 simulations under a choice of parameters), we verified that it was significantly preferred over the runner-up tree to guarantee it was indeed the ‘winning topology,’ and not that it won by sampling variability. PM is obtained by dividing the number of scenarios where the true topology is significantly more likely than the runner-up, by the total number of scenarios considered.

Note that overall accuracy (OA) and proportional mode (PM) may not necessarily be strongly correlated. Suppose, for instance, that for a given model there is a bias in estimation towards certain trees under certain settings. That could result in a large OA because of the large frequency of estimations of the true tree in settings where the model is biased towards it. But if the true tree is not always favored over different simulation settings, the true tree would not be most frequently estimated, leading to a small PM. On the other hand, one could have really low OA but no other tree is chosen more times, leading to a high PM. Ideally, one would like to see both high OA and PM, which would indicate that the true tree is being chosen the most and with little variability across scenarios.

## Results

We present the results according to the set of mixing distributions we used to simulate the data. We start with ℭ60, and then UDM. We conclude the results section by looking at some empirical data sets. These latter are meant to be an exploration of the performance of models for large numbers of sites under misspecification, where the consistency results do not apply.

For the remainder of the text and for simplicity, we denote by C60L the model ℭ60 as defined in Si Quang et al. (2008) including both frequency classes and optimal weights from that study. Then, when we discuss fitting one of the CXX models we are referring strictly to fitting just its frequency classes.

### Simulated Data Under ℭ60 Mixing Distributions

Table 1 displays the mean ISE for data generated under 72 different simulation conditions total; i.e. under eight mixing distributions in ℭ60 and nine trees. The eight mixing distributions included C60L and C60[*i*] for *i* in *{*10, 15, 20, 30, 40, 50, 60*}*. We see in this table that fitting the true generating models with an F-class, denoted here as perfect fit (PF)+F, produces the lowest ISE. The second lowest value is achieved by C60+F model, followed by C40+F, C30+F, and C20+F, respectively. Therefore even though C60+F fits 61 classes to the data – more than the true model – it has a much better fit than the CXX+F models with fewer classes. This suggests that over-parameterization is not a problem for complex, correctly specified models.

**Table 1.**
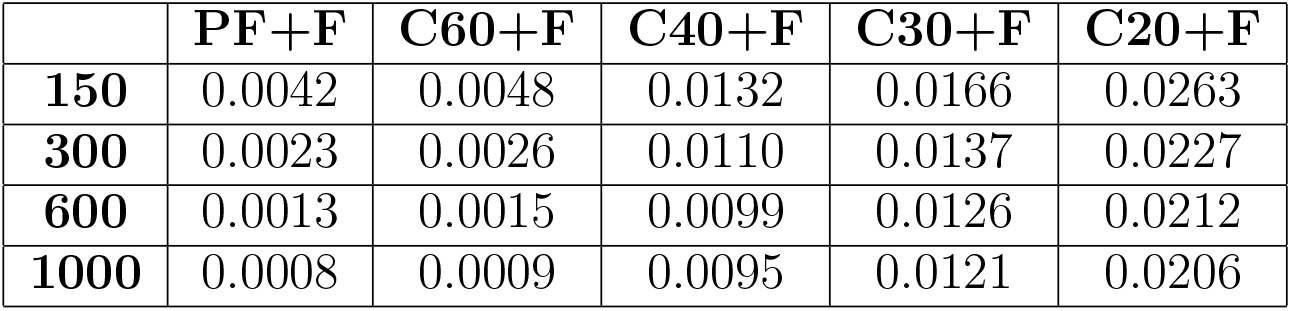
The mean ISE for data generated under distributions in ℭ60. For each of the 72 scenarios (9 trees and 8 models) per sequence length, we compute the mean ISE for the 100 simulations. Then we take the mean of all these values, which are the entries in this table. Label PF+F represents the overall performance in ISE when fitting the generating classes per scenario with the F-class.

Table S2 in the Supplementary Material shows the mean normalized ISE for the same set of simulations, where ISE for any given scenario was re-scaled to give a sum of 1 over all fitted models. This shows that the mean ISE values in Table 1 adequately consolidate all scenarios and no biases between classes are introduced by a scenario with considerably larger ISE values.

Figure 2 shows the plot of average overall accuracy, OA (A), and proportional mode, PM (B), over all data generated under mixing distributions in ℭ60. Each dot in OA represents the proportion of times the tree was recovered over 2400 simulations (3 values of *l* and 8 sets of classes in ℭ60, with 100 repetitions each), and PM is a proportion over 24 distinct scenarios. From PM and for all alignment lengths, we see that models with fewer classes behave worse than ‘richer’ ones. This becomes more prominent as sequence length increases. This aligns with the ISE observations, where complex models that approximate the true CDF better, appear to have superior performance.

**Fig. 2.**
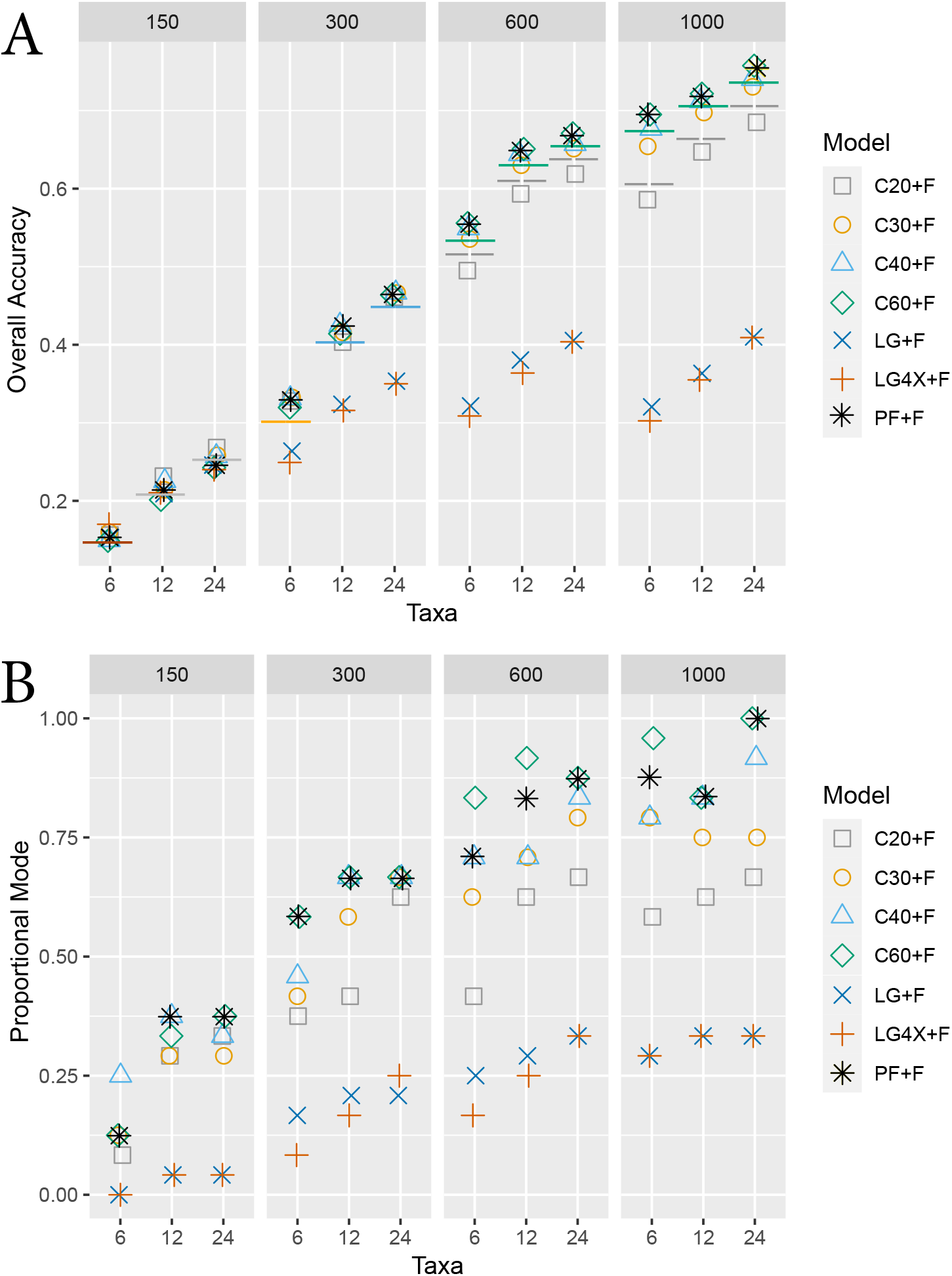
(A) the plot of the overall accuracy (OA) values for various models fitting data generated under mixing distributions in ℭ60 fitted to all frequency vectors. Label PF denotes the frequency vectors used to generate the data. The *x*-axis represents the number of taxa on the tree. The plot is divided by sequence lengths of 150, 300, 600, and 1000. The lower bound of the 95% confidence interval (CI) of the model with the highest OA is depicted with a solid line whose color is in agreement with such model. Depicted with a dashed gray line is the higher bound of the CI of classes in C20 in the cases where such model was significantly worse than the best model. (B) A similar plot to that on top but for proportional mode (PM). For this case, no confidence interval can be computed.

The OA are comparable for very short alignments but this quickly changes as sequence length increases. We see that the simplest models, LG+F and LG4X+F, behave poorly. We also see that for longer alignments, ‘richer’ complex mixture models, like C40+F, outperform C20+F. Overall these results reinforce the inference that: (1) over-parameterization of C60+F does not cause problems; and (2) misspecification of the frequency vectors does not compromise tree estimation if the estimated CDF adequately approximates the true CDF.

One concern is that C60+F could be fitting better on average because it closely resembles two of the mixing distributions – C60L and C60[60] – used in the foregoing simulations. However, C60+F also behaves well for data generated with fewer classes. Table S3 in the Supplementary Material shows the OA for data generated under C60[*i*], with *i* ∈ 10, 30, 60, *T*_6_(0.005), and different sequence lengths. The data in this table indicates that problems with over-fitting do not arise when the estimating model has many more classes than the generating model. This is also reflected in the ISE scores when considering different generating classes (Supplementary Material Table S4). We note that for both OA and PM we see that estimation is improved when there are more taxa present (Table 2), consistent with what we would expect from the ISE scores. This behavior is consistent across models.

**Table 2.** The ISE for fitted C60+F to data generated under *T*_6*m*_(0.02) and C60[15] for all *m*. One can see a decrease in ISE by either increasing the number of taxa, or the sequence length.

| Taxa | ISE |  |  |  |
| --- | --- | --- | --- | --- |
|  | 150 | 300 | 600 | 1000 |
| 6 | 0.00474 | 0.00249 | 0.00124 | 0.00079 |
| 12 | 0.00391 | 0.00187 | 0.00099 | 0.00050 |
| 24 | 0.00279 | 0.00144 | 0.00075 | 0.00039 |

To confirm that LBA is the main source of bias when the true tree is not recovered, Figure S2 shows for this set of simulations the proportion of times the LBA tree is selected and the proportional mode of the LBA tree. It is clear that when the simpler models fail to recover the true tree, the LBA tree is generally recovered.

We finish this subsection by commenting on some supplementary simulations on larger trees (see the complete derivation in the Supplementary Material). The first set of simulations was from a 40-taxon tree sampled from a birth and death process. We simulated alignments as done in this section but using these trees and alignment lengths of 100 and 600. For the simulations obtained from the birth-death process tree, we performed a full tree search on IQ-TREE 2 for all fitted models. Roughly, 78% and 94% of the non-trivial splits in the tree are correctly inferred for alignments of lengths 100 and 600, respectively. These percentages are higher than the OAs in the previous simulations discussed above. We believe this is because (1) most of the splits of the birth-death tree suffer from LBA biases and (2) the greater taxon sampling of in these trees helps improve split inference. For these simulations, we see no significant difference between fitting the true model and C60. Additionally, we see that although ℭ60 only marginally outperforms the slightly less complex mixture models (e.g. C40+F), it does significantly outperform C20+F and simpler models (e.g. LG4X+F and LG+F), and is never outperformed by any other model (Table S5.5). Box plots for the weight estimates for all classes in C60+F (Figure S4) show that the weight estimates are well approximated. This is consistent with the small ISEs for ℭ60 as is expected because of the separation of frequency vectors in the ℭ60 model Further simulations were conducted using the larger trees considered in the real data analyses (I and III) below but with smaller sequence lengths. The overall results of these simulations are presented in Tables S6-S8 and are in good agreement with the findings from the birth and death simulations. The estimated trees from the C60+F and C40+F fitted models gave relatively large proportions of correct splits, comparable to PF+F and significantly better than simpler mixture models (e.g. C20+F, LG4X+F) and those that do not involve profile mixtures. In comparing likelihoods for the true tree and an artefactual LBA tree, C60+F had the largest frequency of preference for the true tree (Table S7). A more detailed discussion of these simulations can be found in the Supplementary material.

### Simulated Data Under UDM Mixing Distributions

Table 3 displays the mean ISE values over data generated under UDM-0256 with POISSON exchangeabilities. The ISE values are considered separately when fitting CXX models (with and without the F-class) to the POISSON and LG matrices. In the former case, there is misspecification just of the classes whereas, for the latter, both the classes and exchangeabilities are misspecified.

**Table 3.** The mean ISE for data generated under UDM-0256 and POISSON exchangeabilities. For each of the 9 scenarios (9 trees) per sequence length, we compute the mean ISE for the 100 simulations. This is done separately when fitting the LG and POISSON matrices.

| ISE | LG matrix |  |  | POISSON matrix |  |  |
| --- | --- | --- | --- | --- | --- | --- |
|  | 300 | 600 | 1000 | 300 | 600 | 1000 |
| **C60+F** | 0.0486 | 0.0529 | 0.0557 | 0.0052 | 0.0041 | 0.0036 |
| **C40+F** | 0.0516 | 0.0557 | 0.0585 | 0.0058 | 0.0046 | 0.0041 |
| **C30+F** | 0.0575 | 0.0612 | 0.0634 | 0.0062 | 0.0051 | 0.0046 |
| **C20+F** | 0.0617 | 0.0652 | 0.0671 | 0.0072 | 0.0061 | 0.0057 |
| **C60** | 0.0232 | 0.0246 | 0.0252 | 0.0050 | 0.0039 | 0.0034 |
| **C40** | 0.0218 | 0.0222 | 0.0226 | 0.0055 | 0.0044 | 0.0039 |
| **C30** | 0.0278 | 0.0289 | 0.0294 | 0.0058 | 0.0048 | 0.0044 |
| **C20** | 0.0261 | 0.0262 | 0.0266 | 0.0067 | 0.0058 | 0.0054 |

When fitting the LG matrix, i.e where there is misspecification of the exchangeabilities, we see a significant difference in model fit between models including the F-class and omitting it. The ISE scores are elevated for models including the F-class relative to those without and this effect is independent of the sequence length (Table 3). We believe this is not directly an artifact of the inclusion of the F-class *per se*, as discussed below.

When fitting the correctly specified POISSON matrix, for any given sequence length and set of fitted frequency vectors, the mean ISE is comparable whether the F-class is included or omitted. In most cases, we see a slightly better ISE when excluding the F-class although the difference is small and most likely not significant. We note that the classes in ℭ60 have the best ISE scores across all sequence lengths, but many of the other CXX models also performed well; C20 yielded the poorest scores overall. In this same table, we also see how ISE decreased as sequence length increased when there is no misspecification of exchangeabilities (i.e. when POISSON exchangeabilities are used to fit). We believe this comes from the fact that as more data becomes available, sites corresponding to rarer UDM classes will be present in increasing numbers giving the richer (and more flexible) higher dimensional mixing distributions a modeling advantage over the simpler models. It is notable that when exchangeabilities are misspecified (e.g. using POISSON to simulate and LG exchangeabilities to fit), however, the richer mixing distributions are still better fitting than simpler ones, but as sequence length increases the ISE generally gets larger for all models. This suggests that misspecified exchangeabilities systematically compromise profile mixture model fitting. Figures 3 and 4 show the plots of average overall accuracy, OA (A) and, proportional mode, PM (B) for data generated under UDM distributions and POISSON exchangeabilities; Figure 3 shows data fitted using models with POISSON exchangeabilities, and Figure 4, LG exchangeabilities. In these plots, each dot in the OA plot represents the proportion of times the tree was correctly inferred over 600 simulations (3 values of *l*, 2 sets of UDM classes, and 100 repetitions), and for PM the proportion over 6 distinct scenarios.

**Fig. 3.**
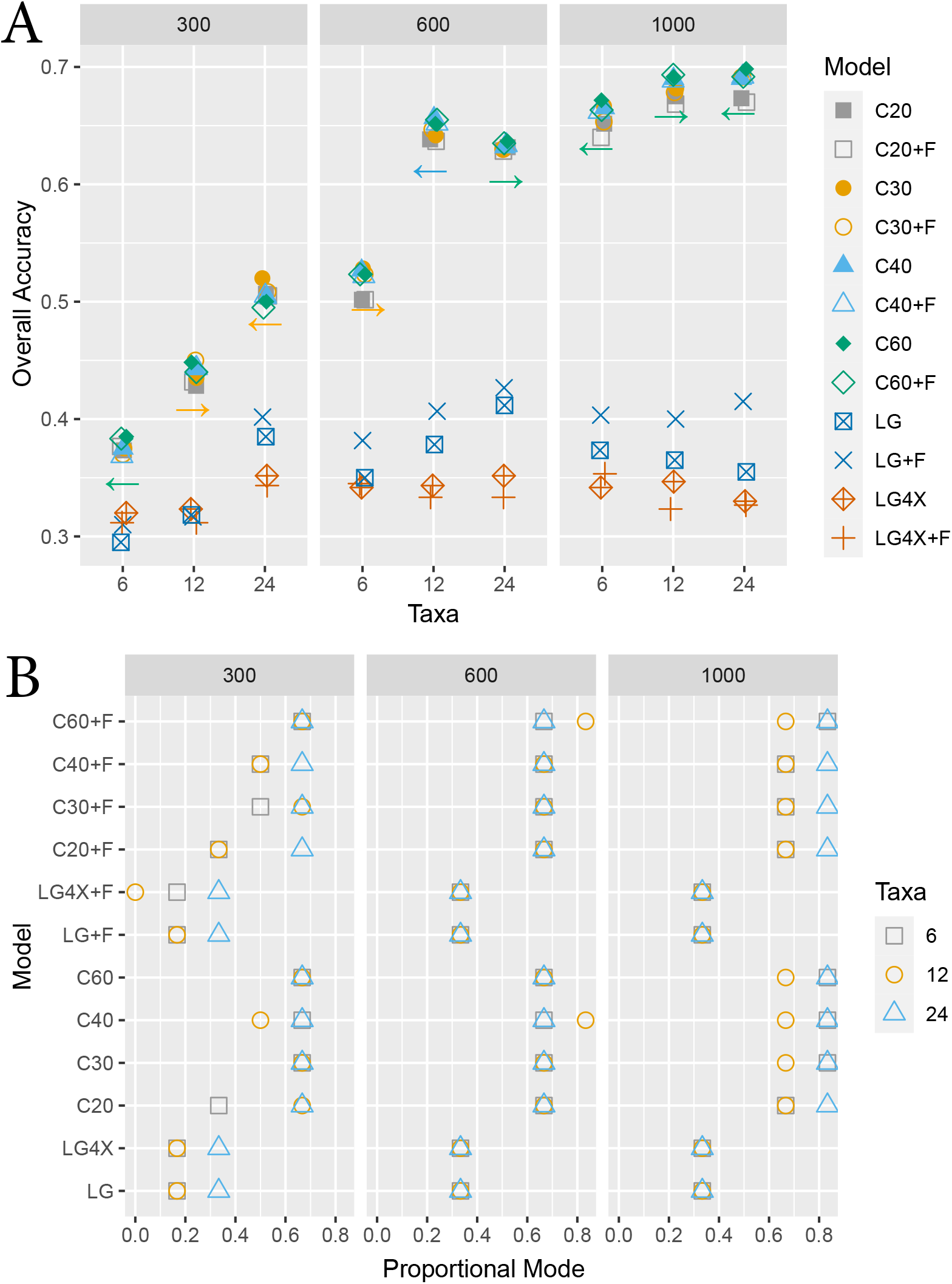
(A) The plot of the overall accuracy (OA) values per model of the data generated under the UDM distributions and POISSON exchangeabilities. Different classes are fitted but in all cases, we fit POISSON exchangeabilities. The *x*-axis represents the number of taxa on the tree. The plot is divided by the sequence length. The lower bound for the 95% confidence interval (CI) of the model with the highest OA is depicted with an arrow whose color represents such model. An arrow pointing to the left represents CXX+F, an arrow pointing to the right represents CXX without F. (B) A similar plot to that on top but for proportional mode (PM). For this case, no confidence interval can be computed.

**Fig. 4.**
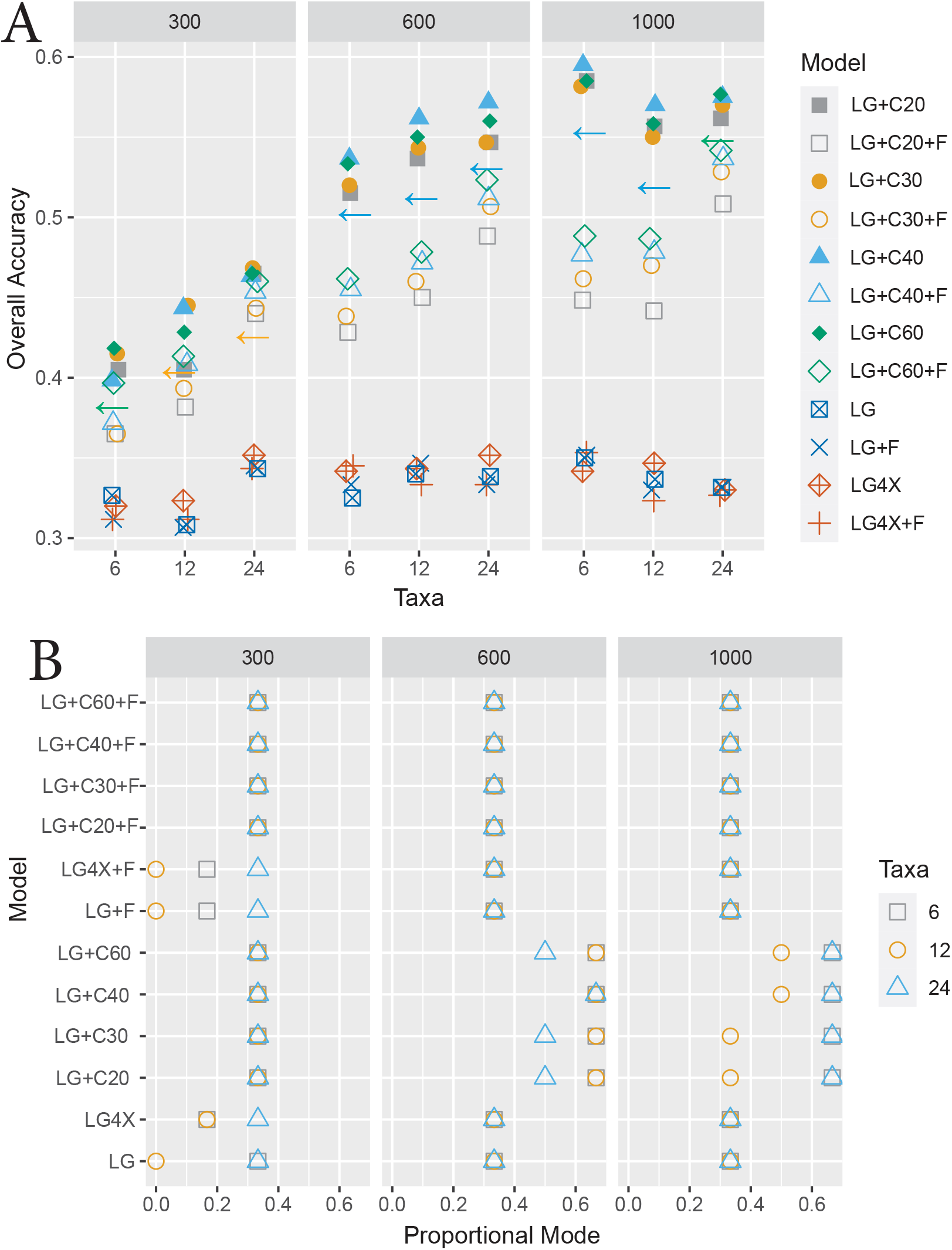
(A) The plot of the overall accuracy (OA) values per model of data generated under the UDM distributions and POISSON exchangeabilities. Different classes are fitted but in all cases, we fit LG exchangeabilities. The *x*-axis represents the number of taxa on the tree. The plot is divided by the sequence length. The lower bound for the 95% confidence interval of the best model per scenario is depicted with a gray line. (B) A similar plot to that on top but for proportional mode (PM). For this case, no confidence interval can be computed.

Figure 3 shows the case when there was misspecification of classes only. Recall that the mixing distributions in UDM have 256 and 4096 frequency vectors, thus there is significant misspecification when fitting CXX models. Nevertheless, we saw that in all cases, fitting with any of the CXX mixtures led to reasonable performance. The OA estimates were even close to the case of no misspecification of the frequency classes in Figure 2.

In this case, there is no significant difference in OA between fitting an F-class as part of the model or not. However, fitting without the F-class yielded, in some cases, better PM values. Similar to the ℭ60 case, fitting with LG4X and POISSON with no site-profile mixture model both behaved very poorly. In this case, we also see how the increase of taxa monotonically improved tree estimation. The exception was for sequence lengths of 600 and 1000 when increasing taxa from 12 to 24. We consider these to reflect a virtual tie in performance with no significant difference between the 12 and 24 taxon simulations. Similar conclusions can be drawn for the case where the only difference was that the data was generated and fitted using the LG matrix instead of POISSON (Supplementary Material Figure S5).

When there is misspecification of both classes and exchangeabilities as shown in Figure 4, we observe that for all cases, except one, OA was equal or significantly worse compared to Figure 3 and Figure S5 in the Supplementary material. We believe this is similar to the behavior shown in Table 3 described above. Furthermore, in some cases, an increase in numbers of taxa led to a decrease in OA (e.g. see Figure 4, going from 6 to 12 taxa with 1000 sites).

The most striking, and perhaps surprising, feature in Figure 4 is that fitting with the F-class often led to a substantial drop in OA compared to when it is omitted. To investigate this further, we explored the impact of the F-class on the likelihood scores and mixing weights of the fitted models when exchangeabilities were misspecified (LG) versus correctly specified (POISSON). These analyses were based on 21600 observations (9 trees, 3 sequence lengths, 2 UDM distributions, 4 fitted models, 100 simulations) and the results are shown in Figure 5. When there is misspecification of the exchangeabilities, the F-class frequently improves the likelihood values substantially, whereas when the exchangeabilities are correctly specified, only modest increases in likelihood are seen (see Figure 5 (A)). The frequently large increases in likelihood values with the F-class in the case of the LG exchangeabilities is surprising because, as indicated in Figure 4 and Table 3, better OA estimates and lower ISE scores are obtained in this case when there is no F-class.

**Fig. 5.**
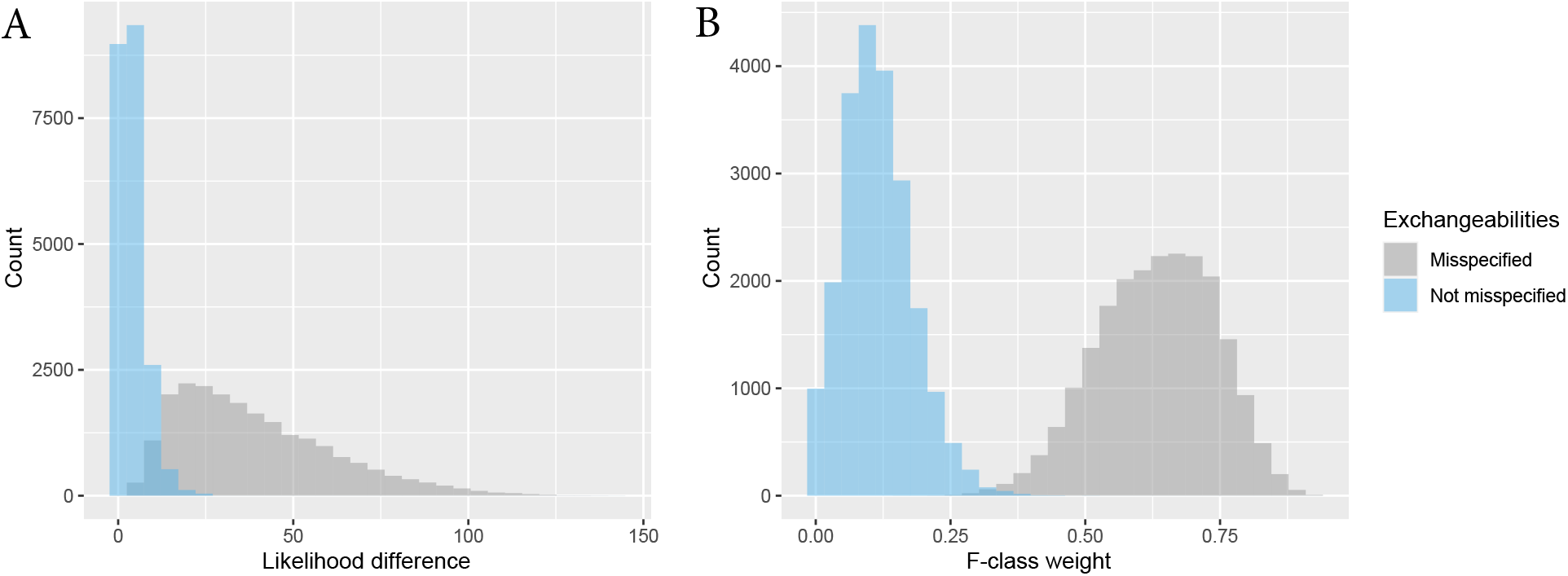
(A) Histograms showing the difference between likelihoods values of models fitted with and without the F-class. No misspecification of the exchangeabilities is depicted in blue and in gray when there is. (B) Histograms showing the inferred F-class weight when there is no misspecification of the exchangeabilities (blue) and when there is (gray). The histograms consist of data generated under all trees, number of taxa, both UDM mixing distributions, and POISSON exchangeabilities. Misspecification of exchangeabilities refers to fitting using LG matrix instead of the POISSON matrix.

Since models without the F-class are special cases of the comparable models that include F-classes, we should always expect an increase in likelihood when fitting the latter models. When there is no misspecification of the frequency classes, a crude approximation is that the null distribution has a mean of 5 and a standard deviation of 5.48 (from a mixture of a degenerate uniform[0] and a *χ*^2^ with 20 degrees of freedom, see Self and Liang (1987)). When there is no misspecification, simulations have a mean of 3.8 and a standard deviation of 3.4, smaller than the natural increase in likelihood, alluded to above, that are expected with increases in the number of parameters estimated. By contrast, with model misspecification, the mean and standard deviation are 34.9 and 19.7, respectively. Thus when there is no misspecification of the exchangeabilities, differences in log-likelihoods are pretty small and would be judged small relative to crude chi-square approximations. By contrast, with misspecification, such differences are very large compared to expectations based on the number of parameters estimated.

We also investigated the impact of misspecification of exchangeabilities on the estimated weight of the F-class (Figure 5B). Figure 5 (B) contains two overlapping plots. When exchangeabilities are correctly specified, the F-class weights tend to be relatively small (e.g. mean weight of non misspecified = 0.11), whereas when they are misspecified the weight distribution shifts dramatically to adopt larger values, often exceeding 0.5 (e.g. mean weight of misspecified = 0.62). Clearly, the weights of the F-class are far from zero in the latter case. We conjectured that the fitted LG exchangeabilities places more weight on more uniform frequencies as a way of compensating for the larger-than-expected variation in amino acids coming from the uniform Poisson exchangeabilities; the LG exchangeabilities are much more heterogeneous. To investigate, we used Shannon entropy as a measure of the uniformity of a frequency vector. We found that mixture classes with the highest entropy get assigned really large weights when simulations are performed with POISSON exchangeabilities and fitted with LG exchangeabilities. Since the F-class is often the class with highest entropy, it gets a high weight under these conditions.

We also explored the effects of misspecification of the exchangeabilities for data generated using the LG matrix but fitted with the POISSON exchangeability matrix. In this case, the tree estimation accuracy is also affected (for eg, OA values are in the range of 0.64 to 0.75 for sequence length of 1000 when it is correctly specified vs. a range of 0.60 to 0.71 when misspecified). Figure S6 in the Supplementary Material shows the average OA (A) and PM (B) for this case. In contrast to the previous misspecification scenario (e.g. Figure 4), the F-class does not seem to hinder tree estimation. Figure S7 in the Supplementary Material, the analog of Figure 5, shows how, in this case, the weight of the F-class is close to zero, and therefore there is no likelihood difference with or without it. Furthermore, as detailed in the Supplementary Material, we did not find the weight of classes to be as strongly correlated with the Shannon entropy, as in the previous case.

Finally, we note that it is formally possible that the generally good performance of CXX models in the foregoing analyses could be related to a tendency of these models to prefer topologies where long branches are apart (i.e. they could have a long-branch repulsion (LBR) bias). To test this, we simulated, without model misspecification, from a topology with long branches together, obtained from the tree in Figure 1 after swapping the clade composed of taxa *c*_1_, …, *c*_*m*_, and *d*_1_, …, *d*_*m*_ together with the edge leading to it and the clade composed of taxa *e*_1_, …, *e*_*m*_ together with the edge leading to it. Consequently, the long branches group together to the exclusion of short branches in this simulating tree. Figure S8 in the Supplementary Material, gives the results for the 12 taxon case for this simulating scenario. On average, the estimates in OA and PM for the more complex CXX models are similar to those found when exploring LBA (Figure 3) indicating no bias in favor or against LBA versus LBR trees when they are true. We note that models with fewer classes tend to have better performance in these cases, suggesting these have a slight LBA bias in favor of the true tree. In contrast, the LG and LG4X models are strongly affected by LBA; i.e., they have notably poor performance under the previously discussed conditions where long branches are apart in the simulating tree and extremely good performance under these conditions where the long branches are together in the true tree. A bias towards either the LBR or LBA topologies is not desirable in general. In this sense the CXX models, especially C30-C60, show little bias in either direction and are clearly better choices under these simulation conditions.

### Real Data

To investigate the impact of the F-class and misspecification on large alignments we analyzed three empirical data sets. These data sets are concatenated super-matrices:

I. a 133-protein data set (24,291 sites *×* 40 taxa) assembled to assess the phylogenetic position of the microsporidia in the tree of eukaryotes (Brinkmann et al. (2005)). The microsporidia are specifically related to Fungi but are sometimes recovered as branching outside of all eukaryotes because of an LBA artifact in which they are attracted to the outgroup archaeal sequences. We consider two trees: the correct tree recovered with the LG+C20+F+G model 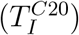 (Susko et al. (2018)) and the LBA tree recovered with the LG+F+G model 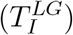 a data set of 146 proteins (35,371 sites *×* 37 taxa) assembled to assess the phylogenetic position of the nematodes in the animal tree of life (Lartillot et al. (2007)). In this case the two competing topologies are the correct topology (recovered with LG+C20+F+G: 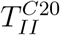) where nematodes branch as sister to arthropods (i.e. the Ecdysozoa group) versus the artifactual topology recovered with LG+F+G 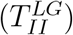.
II. a data set of 146 proteins (35,371 sites *×* 32 taxa) assembled to assess the phylogenetic position of the platyhelminths in the animal tree of life (Lartillot et al. (2007)). The correct position of platyhelminths within the Protostomia is reflected in the tree recovered by CAT+GTR 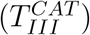 instead of the artifactual Coelomata topology 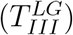 recovered by LG+F+G and many mixture models (see Lartillot et al. (2007), Susko et al. (2018) and Wang et al. (2017))

For each of these trees, we computed the log-likelihoods when fitting classes in C20, C40, and C60, with and without the F-class, and with both LG and POISSON matrices. These likelihoods are shown in Table 4. In all cases, fitting with the LG matrix produces higher likelihood values.

**Table 4.**
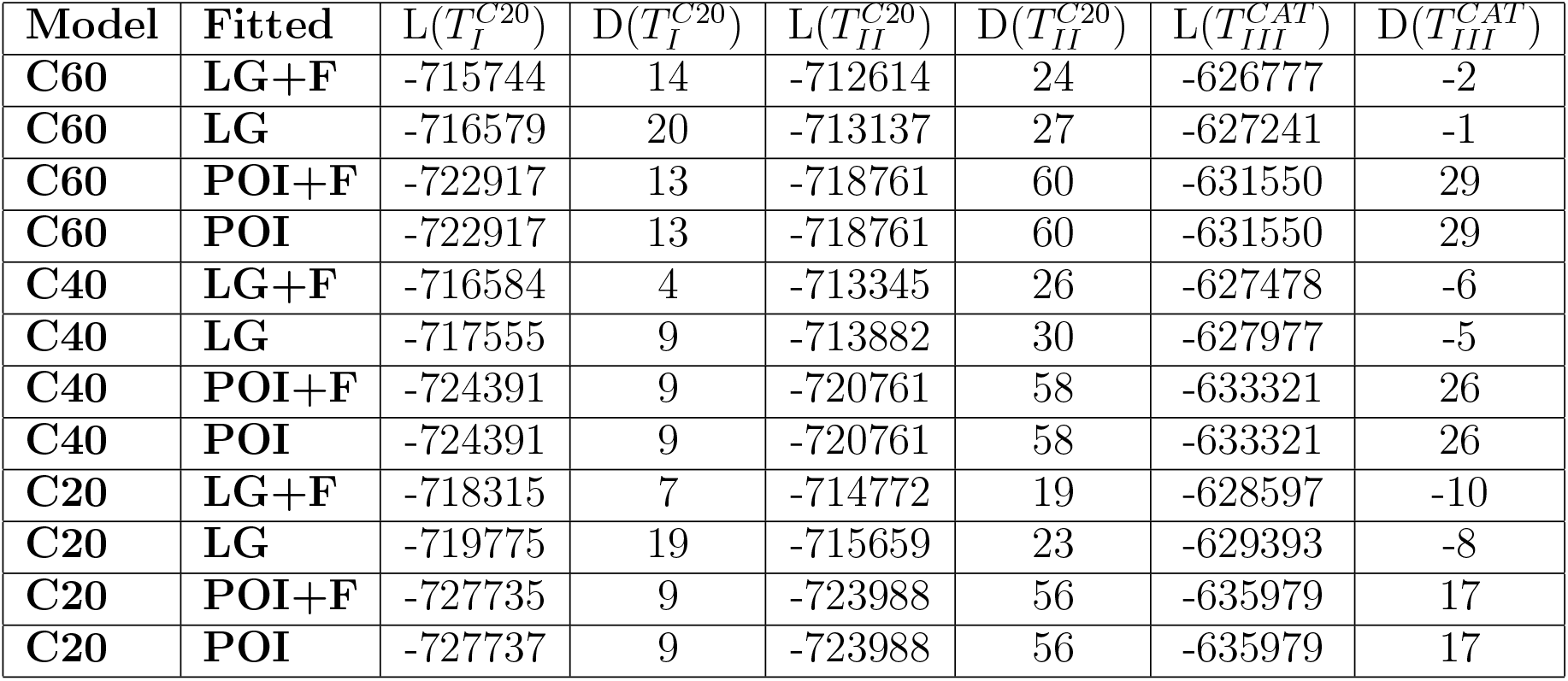
The log-likelihoods of the trees estimated from the empirical data sets, where D(*T*_*J*_) denotes the log-likelihood of the ‘correct tree’ (e.g. C20 or CAT superscripts) minus the ‘incorrect’ tree (e.g. LG superscripts). POI stands for POISSON matrix of exchangeabilities. The best scores are those for C60+LG+F located at the top row.

For data set I, we observe that the correct tree is obtained with the largest log-likelihood differences over the incorrect tree when ℭ60 and C20 are fit with LG exchangeabilities. For this data set the F-class only sometimes negatively affects topological estimation, but never improves it. For data set II, POISSON exchangeabilities strongly favor the correct topology over the incorrect one relative to LG. Here the F-class makes little difference but again never increases support for the correct tree. For data set III, POISSON exchangeabilities favor the correct tree over the incorrect tree, with ℭ60 showing the biggest log-likelihood difference. LG exchangeabilities seem to always favor the incorrect tree.

Overall, the F-class never increases support for the correct tree and sometimes decreases it. Whether LG improves estimation versus POISSON depends on the data set. However, we note that the proteins and taxa in data sets II and III heavily overlap so the outcomes of these analyses are not technically independent.

To conclude this section, we make some comments on additional simulations where we explored misspecification under long alignments. Figure S9 in the Supplementary Material shows the overall accuracy (OA) for simulated alignments of length 20,000 under both UDM distributions, tree *T*_12_(0.005), and different exchangeability matrices. In all cases, there is marked misspecification of the frequency classes. In two of these simulated scenarios, there is also misspecification of the exchangeabilities. In all cases, the simplest models perform even worse than for short alignments. When there is no misspecification of the exchangeabilities, we see clearly how complex models perform better with ℭ60 being consistently the best. For data simulated under POISSON exchangeabilities but fitted with LG, all models recover the wrong tree all the time. For data simulated under LG exchangeabilities but fitted with POISSON, complex models perform better, and again ℭ60 is the best. In none of these does the F-class seem to improve tree estimation. In all cases, richer models performed better. A plot with the proportional mode (PM) values for these simulations is not presented since there are only 2 settings (the two UDM distributions).

## Discussion

Over-parameterization has been an active concern in phylogenetic studies. In a Bayesian framework, the use of complex models versus overly simple models has been strongly favored (Lemmon and Moriarty, 2004; Huelsenbeck and Rannala, 2004). It has been suggested that over-parameterization issues can be avoided since complex models can be regularized through the use of well-behaved priors (see for example the discussion in Guimarães-Fabreti and Höhna (2022)). In a maximum likelihood framework the potential for problems with over-parameterization have been mentioned in many contexts including substitution model selection (Kelchner and Thomas, 2007; Abadi et al., 2019), model adequacy (A Shepherd and Klaere, 2018), partitioned analysis (Seo and Thorne, 2018), and as a potential pitfall in analyses with complex models (Crotty et al., 2019). However, the specific effects of over-parameterization of various kinds of models in ML estimation, has not, to our knowledge, been systematically examined in detailed simulation studies. As mentioned in Sullivan and Joyce (2005), the model with the best maximum likelihood score is not guaranteed to produce accurate estimates from finite data. Increases of likelihoods can also be the result of: (1) indirect effects of parameterizing stochastic variation; or (2) parameters of highly complex models accommodating idiosyncratic patterns in the data coming from other un-modeled processes (see for example (Jones et al., 2018)). Although increased variance in parameter estimates associated with over-fitting is often mentioned (Steel, 2005; Kelchner and Thomas, 2007; Nascimento et al., 2017), truly pathological over-parameterization effects have also been reported such as the inconsistency of the no common mechanism model (Steel, 2010), and the compounded small sample bias associated with over-partitioning data sets (Wang et al., 2019). Here, we have focused on potential problems with over-parameterization in the context of site-profile mixture models of protein sequence evolution. We argue that mixture models have inherently different behavior with respect to over-parameterization than other types of phylogenetic models and parameters. By extending earlier results (Kiefer and Wolfowitz, 1956; Lindsay, 1983), we show that, for profile mixture models, the tree and mixing parameters of profile mixture models are statistically consistent even with a large number of classes. However, since good performance is not guaranteed for short alignments, we conducted an extensive simulation study of the performance and properties of profile mixture models with smaller data sets. We also investigated the effects of model misspecification through both misspecification of the frequency classes and the exchangeabilities for short and long alignments and considered several empirical examples of long phylogenomic-scale alignments. Finally, the effects of an additional F-class were also investigated in all possible settings. These analyses provide useful theoretical and practical insights regarding model fit. Our main findings are the following:

### (A) Over-parameterization is not a problem for complex models

For all alignment sizes explored here, we saw no evidence (in terms of ISE, OA, and PM) that use of more complex profile mixture models led to more variable or poorer estimation. This is true even for models having several classes with zero weights estimates. Consistent with the theoretical results for large numbers of sites, over-parameterization of these mixture models does not appear to be a problem for shorter alignments. Thus, the arguments in Anderson and Lindgren (2021), Li et al. (2021), and Al Jewari and Baldauf (2022) regarding the choice of simpler models to avoid over-fitting should be reconsidered.

Since it is the mixture structure that is important in assessing whether models are over-parameterized, the large sample results we discuss here likely extend to rates-across-sites mixtures (Yang (1994); Felsenstein and Churchill (1996); Mayrose et al. (2005); Susko et al. (2003)) and the codon model-based mixtures used to infer selection pressure (Yang et al. (2000)). We also speculate that some of the small sample results found here may extend to those settings too but additional work is needed to test that prediction.

### (B) Misspecification of the frequency vectors does not necessarily imply bad fit

Misspecification of the frequency vectors in profile mixture models does not cause problems if the estimated CDF can adequately approximate the true CDF. The more data available, the more classes are likely needed to closely approximate the true CDF. As mentioned before, this suggests that many-class models in the CXX (Si Quang et al., 2008) and UDM (Schrempf et al., 2020) families will be best suited for analysis of diverse data sets, assuming computation time is not limiting.

### (C) Simple models behave poorly

Likely as a consequence of (B), both the site-homogeneous POISSON and LG models, and the site-heterogeneous LG4X model perform very poorly in all scenarios. We believe this is because these have one (LG and POISSON) or very few (LG4X) classes. Although inference using simple models can be much faster we do not recommend their use given their poor performance (i.e. susceptibility to LBA) under realistic site-heterogeneous simulation conditions.

Al Jewari and Baldauf (2022) argue that estimated topologies from the LG+Γ model should be trusted over results from the complex site-profile mixture models due to the agreement in tree topology estimates obtained from the former model when fitting differently filtered data sets. Our results suggest that this observation could be attributed to strong biases in the LG+Γ-based estimates that are consistent over data sets rather than to better performance.

### (D) Misspecification of exchangeabilities and the presence of an F-class can severely affect tree estimation

A severe decrease in accuracy of tree estimation is observed for data generated under the POISSON exchangeability matrix and fitted using the LG exchangeability matrix when an F-class is included in the mixing distribution. In these scenarios, it is clear that the F-class has large estimated weights and degrade performance of the model. For data generated under the LG matrix and fitted using the POISSON matrix, performance is affected but not as severely as in the previous case.

We suggest that misspecification of exchangeabilities can be problematic when the matrix used to fit the data has less uniform exchangeabilities than the true matrix. The exchange rates for empirical rate matrices like the LG matrix were estimated from large databases using likelihood approaches that did not include profile mixtures. It is likely that some of the site-specific amino acid preferences were, under these conditions, accommodated by increased exchangeabilities between these amino acids in the LG matrix. We therefore suspect that estimates of exchangeabilities would be more uniform if profile mixtures models like CXX were used in the estimation of the exchangeability matrix. The poor performance associated with an F-class may apply to real estimation settings. We note that the use of both the LG exchangeability matrix and the F-class leads to higher likelihoods compared to POISSON exchangeabilities and no-F-class, so model selection criteria like AIC will frequently favor their use in real settings. Since the F-class never appeared to improve estimation in any of the simulations or real data analysis settings we examined, we discourage its use in site-profile mixture models.

### (E) Better likelihood scores do not imply better tree estimates

As a consequence of (D), and also observed in the empirical data analyses, higher log-likelihood scores do not always imply better tree estimates. This is also weakly observed even when there is no misspecification of the exchangeabilities.

### (F) Adding more taxa can improve or hamper tree estimation accuracy

Adding more taxa generally improves ISE and tree estimation. Surprisingly, when the model is misspecified (e.g. using UDM frequencies and misspecification of exchangeabilities) adding taxa does not always improve estimation; in one case it actually decreases performance (Fig. 4).

The poor performance of the methods when exchangeabilities are misspecified provides a strong motivation to develop software tools that allow ML estimation of a GTR matrix over all sites in the presence of a profile mixture model. In future work, we plan to construct mixing distributions that closely approximate the true CDF for data, hoping this would lead to a more accurate tree estimation than current models. One could argue that relying on a single set of exchangeabilities being shared across sites with differing frequency classes may be an artificial construct. Nonetheless, we hope that even if it is the case, it would still perform better than using an *a priori* empirically derived matrix.

As mentioned, misspecification of the frequency vectors in profile mixture models does not cause problems if the estimated CDF can adequately approximate the true CDF. For a mixture model to be able to approximate the true CDF, the frequency classes must be in the neighborhood of the classes of the true model. In reality, having a large number of classes is not necessarily enough to achieve this, since, in principle, all classes could be far away from the truth. Larger models in the CXX or UDM families perform, in general, adequate because they were estimated on large databases of alignments and therefore capture commonly occurring site profiles. But even if such models have extra classes not represented in the particular data set of interest, it does not severely affect tree topology estimation and the ML estimates of their weights will often be close to zero. In general, we recommend the use of class-rich models which are likely to better approximate the CDF of the true mixing distribution. Thus, the only remaining downside of fitting very rich profile mixture models, is the significant computational burden they impose on calculation of log-likelihoods for large datasets.

From the theoretical framework in Kiefer and Wolfowitz (1956) and Lindsay (1983) the results on long alignments presented here can be easily extended to other mixture contexts. For example, one could easily extend these results to an analogous of profile mixture models but on a nucleotide or codon setting. For short alignments, the implications are not as clear, since both nucleotide and codon models could be troubled by saturation and compositional bias issues at some sites (e.g. at nearly-neutral sites such as third codon positions in protein genes). Although site profile mixture models do discard information because they do not directly model the fundamental molecular evolutionary process of nucleotide substitution, the saturation and compositional bias issues are less likely to be present.

Finally, the foregoing results lead to some practical recommendations for single and multi-gene phylogeny inference. First, we do not discourage the use of ‘rich models’ (those with many frequency classes), even when several classes have zero weight estimates. Conversely, we do suggest avoiding models with one or very few frequency classes. We also discourage the use of the F-class, unless analyses are done both with and without the F-class, and the results are critically examined; if the weight on the F-class is substantial (i.e. *>* 0.25), then the F-class estimates should be treated with caution. Also, it is also worth conducting site profile mixture model analyses both with LG (or other empirical exchangeability matrices) and with the POISSON exchangeabilities to cross-check the sensitivity of the estimation to these settings.

## Appendix

### Cumulative Density Functions and Profile Mixture Models Re-parameterization

The cumulative density function *F*_*X*_(*x*) of a random vector *X*, evaluated at *x* is the probability that *X* takes values less or equal to *x*, that is *F*_*X*_(*x*) = *P* (*X* ⩽ *x*). When *X* = (*X*_1_, …, *X*_*q*_) is a multivariate random vector, the cumulative distribution is defined as the joint probability that *X* takes values less or equal to *x* = (*x*_1_, …, *x*_*q*_), that is

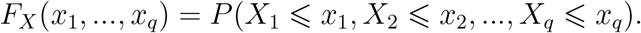

We now tie these concepts to our particular scenario. For simplicity, we do this first in the nucleotide setting (dimension 4), rather than the amino acid setting (dimension 20) but all ideas extend naturally to the latter setting. The cumulative density function of the nucleotide frequency at a site is given by:

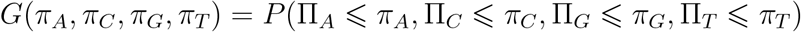

where (Π_*A*_, Π_*C*_, Π_*G*_, Π_*T*_), and Π_*i*_ denotes the random frequency of nucleotide *i* at a site. Thus if ***π*** = (0.5, 0.3, 0.4, 0.7)

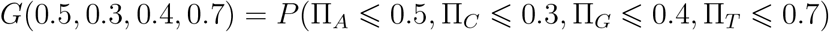

denotes the joint probability that the frequency of nucleotides (A,C,G,T) is less or equal than (0.5, 0.3, 0.4, 0.7), respectively, at a site.

We now exemplify how cumulative distributions can be constructed in the context of profile mixture models. We do this in the simpler setting of purines and pyrimidines (RY-coding), ***π*** = (*π*_*R*_, *π*_*Y*_), for a mixture with the following 3 frequency classes:

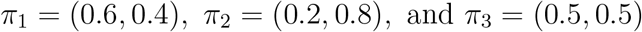

with mixture weights *w*_1_ = 0.1, *w*_2_ = 0.4, and *w*_3_ = 0.5. Then

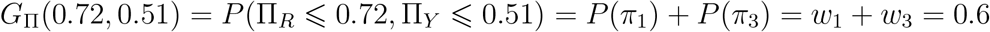

This is because in both frequency vectors *π*_1_ and *π*_3_, the frequency of *R* is less or equal to 0.72 and the frequency of *Y* is is less or equal to 0.51. The probability of this event is the sum of the probabilities of each frequency vector (given these events are disjoint). Figure 6 shows the plot of the contour lines of *G*_Π_(*π*) for this example.

**Fig. 6.**
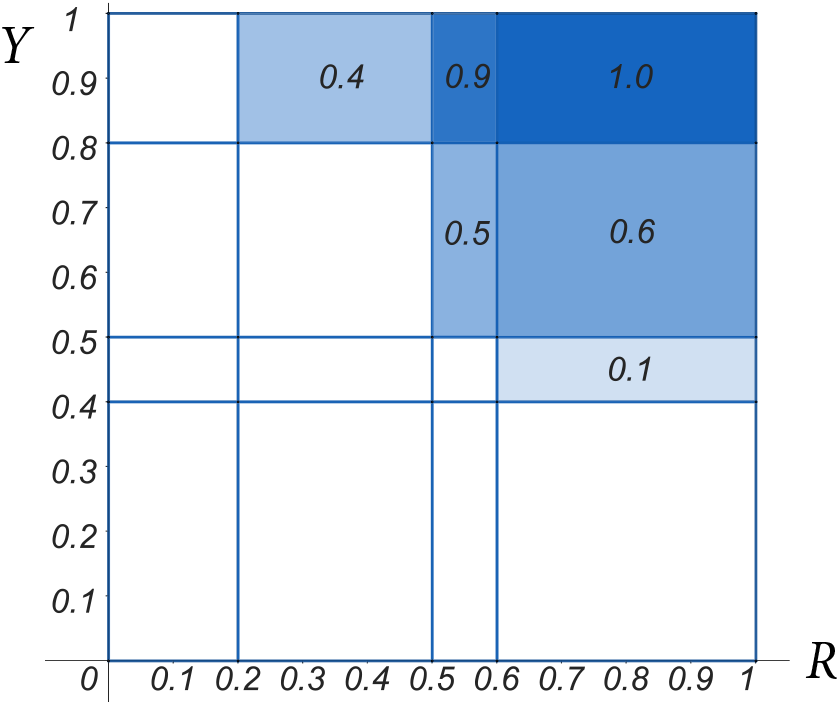
The plot of the contour lines of the CDF under the mixture defined by *π*_1_, *π*_2_, and *π*_3_. The x-axis represents the purine (R) frequency and the y-axis the pyrimidine (Y) frequency. Unlabeled regions have zero density.

One advantage of parameterizing in terms of CDFs is that similar profile mixing distributions will have similar CDFs but might have very different *w*_*c*_ parameters or include some very different frequency vectors with small weights. As an example, consider the distribution above but where the *π*_3_ vector is split into two very similar vectors, *π*_3_ = (0.499, 0.501) and *π*_4_ = (0.501, 0.499) with weights *w*_3_ = *w*_4_ = 0.25. These are similar distributions and have similar CDFs but the *w*_*c*_ parameters are not comparable.

#### Statistical consistency of the MLE

As mentioned in the section entitled “*Mixture models and over-parameterization with large samples*”, the results of Kiefer and Wolfowitz (1956) imply that the tree and mixing parameters are consistent. In this section we elaborate on why this is true in the context of site-profile mixture models. We assume the space of mixing distributions is constrained enough that identifiability holds in what follows; identifiability is discussed in the next subsection. The main additional condition that requires argument is the compactification or extension-of-continuity *Assumption 2* in (Kiefer and Wolfowitz, 1956). The rest of the conditions required by Kiefer and Wolfowitz (1956) hold almost immediately upon showing *Assumption 2* for profile mixtures. This is because the probability distributions are continuous functions of parameters and are distributions over a finite, albeit large, space of all site patterns. The latter ensures that expectations of interest are finite sums and always exist.

To deal with the compactification condition, we first introduce the concept of a *forest*. Probabilities on trees that involve infinite edge-lengths, which we need to consider to establish the regularity conditions of (Kiefer and Wolfowitz, 1956), can be expressed as a product of probabilities over phylogenetic trees.

A phylogenetic *forest F* on *X* is a collection of phylogenetic trees *F*_*q*_ on *X*_*q*_, known as *components*, where *X* = ∪*X*_*q*_ and *X*_*q*_ ∩ *X*_*p*_ = ∅ for any two components.

Given the value of (*π*_*c*_, *r*_*k*_) at a site, a site pattern probability for a tree that has infinite edge-lengths can be expressed as a product of probabilities over components of a random forest. To see this, note that every edge in a tree separates the data in two subsets of taxa, *X*_*q*_ and *X*_*p*_, that have no taxa in common. When the edge-lengths are infinite, the data for *X*_*q*_ and *X*_*p*_ are conditionally independent. We can iterate this process to construct a partition of the taxa, by successively partitioning taxa within partitions, whenever there is an infinite edge-length in the subtree, *F*_*q*_, for the taxa *X*_*q*_. It is not difficult to see that the resulting partition, 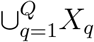 is unique (see for example the split equivalence theorem). So probabilities on trees that include infinite edge-lengths can be expressed as a product of probabilities over the components, each having only finite edge-lengths, of the random forest constructed above. Thus, to deal with infinite edge-lengths, the extension of the space of trees that needs to be considered is

(i’) A metric forest *F* on *N* taxa obtained after removing a, possibly empty, set of edges from a rooted metric binary tree on *N* taxa and retaining only components that display taxa.

The site likelihood contribution then becomes:

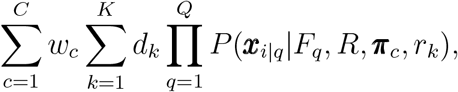

where *Q* are the number of components of *F*, and *x*_*i*|*q*_ is the *i*-th site pattern restricted to the taxa in *F*_*q*_. The reasoning behind forests is to account for infinite edge lengths, which are represented by the edges missing from the tree defining *F*. This not only allows us to consider this limiting case but also, it weakly depicts the effects of functional divergence (Gaston et al. (2011)). Note that when no edges are removed from the tree in (i’), the resulting forest has one component and the likelihood is the same as the one defined in the section “*Mixture models and over-parameterization with large samples*.”

Kiefer and Wolfowitz (1956) decompose the original parameter space (the one before adding on the mixing distribution) into Ω*×* Γ where Γ gives the set of parameters that a mixing distribution will be applied to. In our case Γ is the set of possible (*π*_*c*_, *r*_*k*_) parameters of Δ_19_ *×* ℝ where Δ_19_ is the 20-dimensional unit simplex. The space Ωis then the space of parameters that are fixed across sites, the space of (*F*_*q*_, *R*) in our setting. Although R was not estimated in our examples, the result is slightly more general if we include it.

Consider transformations that convert Ωand Γ to compact (closed and bounded) sets. For *π* and the space of *R*, no re-parameterization is required. Because *R* only needs to be specified up to re-scaling, the space of *R* can without loss of generality taken to be the unit simplex. The spaces that require transformations are the space of *F*_*q*_ and the space of rates. We deal with the latter by re-parameterizing rates via the logistic transformation *ζ* = *e*^*r*^*/*(1 + *e*^*r*^) and similarly re-parameterize edge-lengths as *p* = *e*^*t*^*/*(1 + *e*^*t*^). After transformation, rates and edge-lengths are in [1*/*2, 1). If we can include the parameters *ζ* = 1 and *p* = 1, then the space of rates, after transformation, is compact. Similarly, the space of *F*_*q*_ can now be viewed as a collection of (2*n* − 3)!! closed cubes corresponding the all different tree topologies.

The construction above gives a new space, 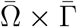, that includes additional parameters, those corresponding to *ζ* = 1 and *p* = 1. The pattern probabilities can be extended to accommodate these parameters by considering the limiting pattern probabilities as *ζ* and/or some of the *p* converge upon 1. These limiting pattern probabilities correspond to cases where rates and/or some subset of edge-lengths become infinite. The concern is then whether they are well defined: Do we get the same pattern probabilities no matter how we approach these parameters? It is not difficult to see that we do. If the limiting *ζ* = 1 (infinite rate) then the model for the patterns independently assigns amino acids to taxa. If the limiting *ζ <* 1 so that subset of the *p* = 1 (some subset of the edge-lengths are infinite), this corresponds to Π_*q*_ *P* (*x*_*i*|*q*_|*F*_*q*_, *R, π, r*) for the forest *F* constructed as described above.

The result of Kiefer and Wolfowitz (1956) applies with constraints on what type of mixing distribution is estimated as long as the constrained spaces of distribution functions are closed. In usual profile mixtures that includes some bound on the number of components considered and that rates are assigned independently from frequency profiles. It is not hard to see that a sequence of distributions satisfying these properties will converge on a distribution that also satisfied these properties. The discrete gamma rate variation model is frequently used. Each rate has equal probability of occurrence and the rates are continuous functions *r*_*k*_(*α*) of a shape parameter *α*. Estimation of the *α* parameter is often constrained to be above a positive lower bound (partly to avoid numerical difficulties) and *r*_*k*_(*α*) → 1 as *α* → ∞, so that the gamma model can be defined on [*α*_0_, ∞] by taking *r*_*k*_(∞) = 1. Consequently, a sequence of rate distributions from a discrete gamma model will converge upon a distribution from a discrete gamma model

#### Identifiability

In this section we argue why, as mentioned in the section “*Mixture models and over-parameterization with large samples*,” for many of the cases considered here the tree parameter is identifiable.

In Theorem 5.7 in Yourdkhani et al. (2021) it is shown that for profile mixture models with *C* · *K <* 72, where *C* is the number of classes and *K* the number of rates, and more than 8 taxa, the tree and numerical parameters are generically identifiable, up to arbitrary re-scaling of the tree and the exchangeability matrix. Generically identifiable means identifiable except maybe in a set of measure zero; informally speaking this means identifiable except maybe in a tiny subset of parameters relative to the full parameter space. Although in such work there is no description of the generic setting of the parameter space, we argue that with just a small perturbation of the parameters one can always guarantee the result to hold.

Since in all models considered here fixed parameters are obtained empirically (that is these have no structure for eg. to be solutions of a phylogenetic invariant) and the parametric function is continuous, there exists ***E*** such that a translation of the numerical parameters by ***E*** will make these generic. This is true for the frequency vectors, weights, rate parameters, edge lengths, and the exchangeabilities matrices. While the POISSON matrix is not generated from data, the proof of Theorem 5.7 in Yourdkhani et al. (2021) is built on this matrix up to a constant and therefore identifiability also holds in this case.

#### ISE Derivation

It is natural to compare two mixing distributions, *H* and *G* (for example estimated and true), by comparing their frequency classes and weights. But this has a number of difficulties including a labeling problem and that similar probability distributions can correspond to seemingly different frequency vectors and/or weights. If the frequency vectors differ, it can sometimes be difficult to determine which frequency class in *H* best matches a given frequency class in *G*. This can be particularly problematic when the numbers of frequency vectors differ substantially between the two distributions. An estimated distribution can also be very similar to the true distribution but with substantial differences in weights and or frequency vectors. For instance, if a frequency vector with weight 1/2 in *H* is approximately the same as two frequency vectors in *G*, each having weight 1/4, *H* and *G* might be very similar but their weights will not be. Similarly, if *G* places small weight on a frequency vector that does not match any frequency vector of *H* well, then the distributions might be similar although the frequency vectors are not. Comparing cumulative distribution functions via ISE(*H, G*) gets around these difficulties.

While in a general setting ISE could be very expensive to compute, we show that in our context (finite mixture of discrete distributions) ISE can be easily computed.

Let ℳ_*s*_ be a profile mixture model, for *s* ∈ *{H, G}*. Let *C*_*s*_ be the number of frequency classes of ℳ_*s*_ for *s* ∈ *{H, G}* and *H* and *G* be the distribution functions of the amino acid frequencies at a site of ℳ_*H*_ and ℳ_*G*_, respectively. For notation simplicity, let *H*_*bi*_ and *G*_*bi*_ be the frequency of amino acid *b* in the *i*-th class for models *M*_*H*_ and *M*_*G*_, respectively. Let *h*_*i*_ and *g*_*i*_ be the weights of frequency class *i* in model ℳ_*H*_ and ℳ_*G*_, respectively.

Note that for *x* = (*x*_1_, …, *x*_*p*_) and *D* = [0, 1]^*p*^

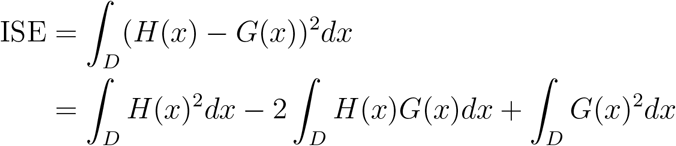

Note that, in particular,

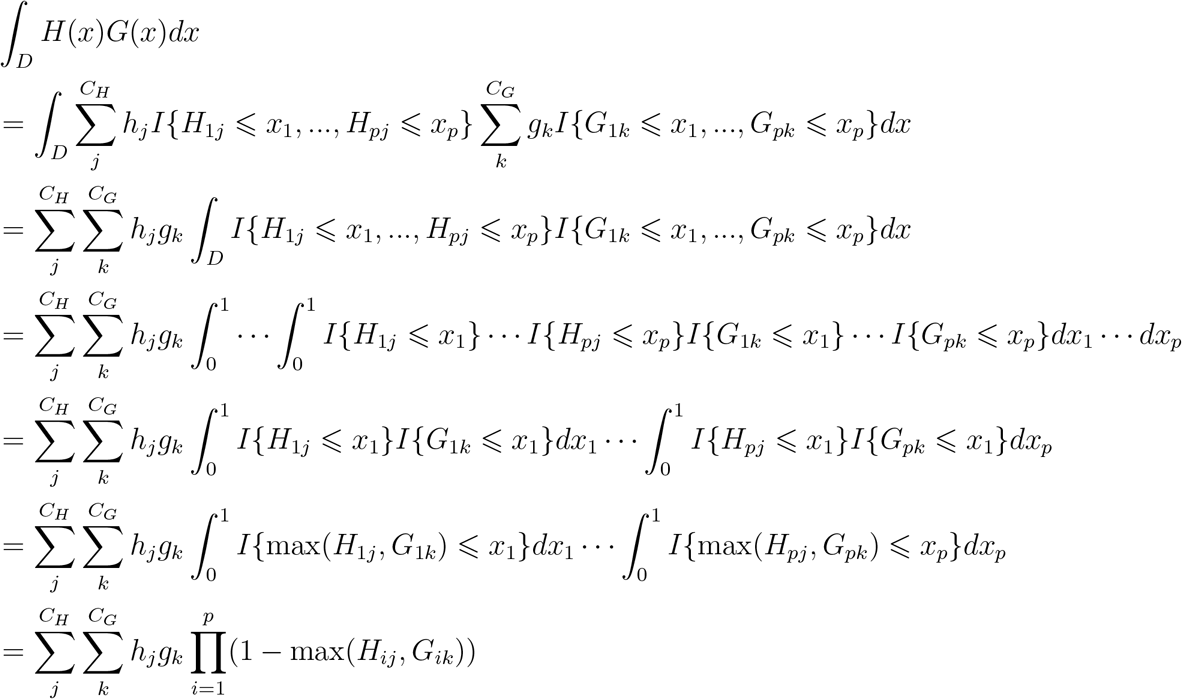

By doing similar computations for the other two terms we obtain:

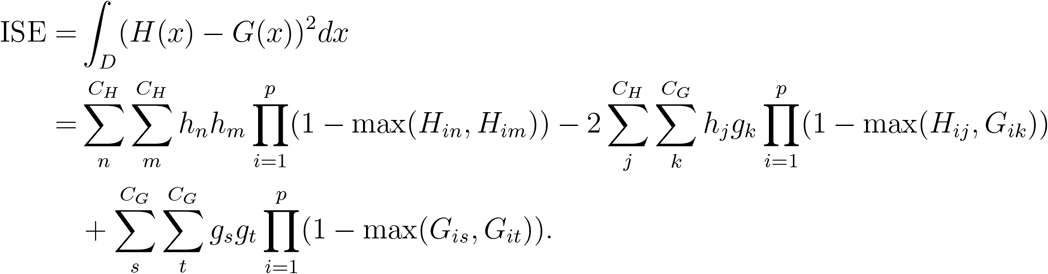

