## Supplementary material for "Is Over-parameterization a Problem for Profile Mixture Models?"

### *Supplementary Figures, Tables, and Sections*

All the materials are presented in the order they are referred to in the manuscript. At the end of this document, we include the section entitled *Reproducibility*, which explains some of the simulation procedure, as well as some sample command lines like used in this work to simulate and fit the data. Throughout this material, blank spaces are left on purpose.

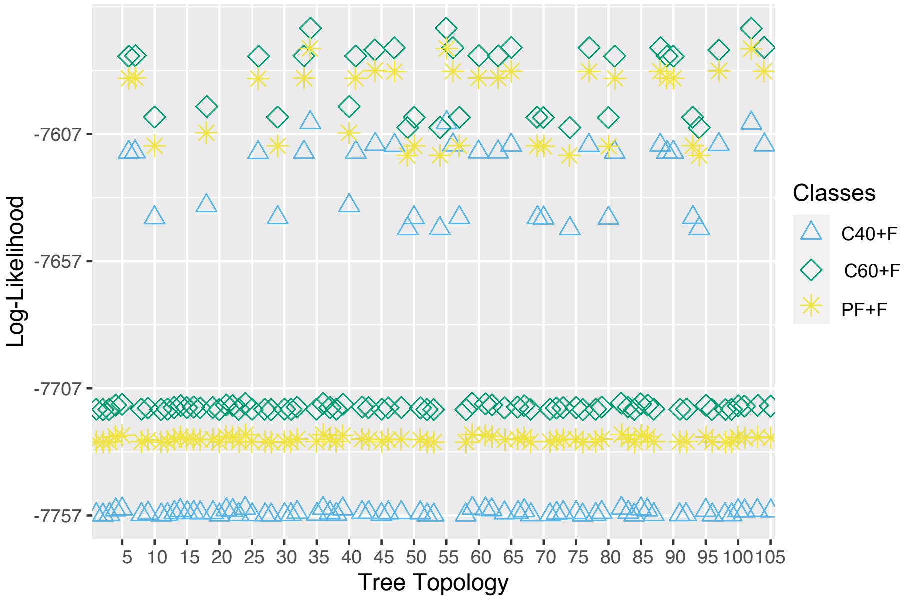


Fig. S1. The log-likelihood obtained from fitting different models from simulated data under $T_6(0.005)$, C60[10], sequence length of 1000, and LG exchangeabilities. The $x$-axis represents the distinct 105 unrooted topology on 6-taxa. The 35 topologies with higher likelihood are those in Table S1 which are the topologies displaying the quartet $AB|CD$.

| **The 35 unrooted 6-taxon topologies displaying the quartet AB\|CD** | | | |
|---|---|---|---|
| (C,D,(F,(E,(A,B)))); | (D,(F,C),(E,(B,A))); | (E,(F,A),(B,(D,C))); | (A,(E,B),(F,(D,C))); |
| (D,C,((A,B),(F,E))); | (C,D,(A,(B,(E,F)))); | (A,B,(C,(E,(F,D)))); | (D,C,(B,(F,(A,E)))); |
| (F,(A,B),(E,(D,C))); | (A,(E,F),(B,(D,C))); | (B,A,((C,F),(D,E))); | (B,E,(A,(C,(F,D)))); |
| (A,(B,F),(E,(D,C))); | (D,(F,E),(C,(A,B))); | (F,(E,D),(C,(A,B))); | (B,A,(D,(F,(E,C)))); |
| (F,(D,C),(B,(E,A))); | (D,C,((A,E),(B,F))); | (D,F,((A,B),(C,E))); | (A,E,(B,(C,(D,F)))); |
| (E,(D,C),(B,(A,F))); | (F,E,(C,(D,(A,B)))); | (B,F,(A,(C,(E,D)))); | (B,(F,A),(D,(C,E))); |
| (E,(B,A),(C,(F,D))); | (E,(B,F),(A,(D,C))); | (F,B,(A,(D,(C,E)))); | (B,(E,A),(D,(F,C))); |
| (A,B,(F,(C,(E,D)))); | (A,F,((D,C),(B,E))); | (B,(F,A),(C,(D,E))); | (A,(E,B),(D,(F,C))); |
| (F,(B,A),(D,(E,C))); | (A,(D,C),(F,(B,E))); | (B,A,(D,(E,(C,F)))); | |

Table S1. The 35 topologies belonging to the tier with higher likelihoods as discussed in the Section *"Fitted models and Precision of parameter estimation"*, and exemplified in Figure S1. All these topologies display the quartet $AB|CD$.

| ISE | PF+F | C60+F | C40+F | C30+F | C20+F |
|---|---|---|---|---|---|
| **150** | 0.0731 | 0.0793 | 0.1866 | 0.2292 | 0.3493 |
| **300** | 0.0615 | 0.0670 | 0.2095 | 0.2590 | 0.4030 |
| **600** | 0.0440 | 0.0475 | 0.2135 | 0.2690 | 0.4260 |
| **1000** | 0.0326 | 0.0353 | 0.2158 | 0.2756 | 0.4408 |

Table S2. The mean normalized ISE for data generated under distributions in $\mathfrak{C}60$. For each of the 72 scenarios (9 trees and 8 models) per sequence length, we compute the mean ISE for the 100 simulations. Then we divide the average ISE of each fitted model by the sum of the average ISE over all fitted models. Then we take the mean of all these normalized values, which are the entries in this table. Label PF+F represents the overall performance in ISE when fitting the generating classes per scenario with the F-class. Values can be compared only across columns, not rows.

| | C60[10] | | | C60[30] | | | C60[60] | | |
|---|---|---|---|---|---|---|---|---|---|
| **Model** | **300** | **600** | **1000** | **300** | **600** | **1000** | **300** | **600** | **1000** |
| PF | 0.25 | 0.39 | 0.46 | 0.24 | 0.41 | 0.41 | 0.21 | 0.39 | 0.43 |
| C60 | 0.26 | 0.43 | 0.48 | 0.26 | 0.43 | 0.44 | 0.21 | 0.39 | 0.43 |
| C40 | 0.22 | 0.41 | 0.49 | 0.24 | 0.38 | 0.39 | 0.23 | 0.39 | 0.38 |
| C30 | 0.19 | 0.42 | 0.49 | 0.22 | 0.34 | 0.40 | 0.24 | 0.38 | 0.39 |
| C20 | 0.17 | 0.31 | 0.36 | 0.22 | 0.32 | 0.32 | 0.29 | 0.37 | 0.32 |
| LG4X | 0.08 | 0.14 | 0.13 | 0.17 | 0.13 | 0.13 | 0.11 | 0.04 | 0.01 |
| LG | 0.11 | 0.20 | 0.16 | 0.18 | 0.14 | 0.18 | 0.10 | 0.06 | 0.03 |

Table S3. The OA values for data generated under $T_6(0.005)$ for all different sequence lengths, and models C60[$i$], for $i \in \{10, 30, 60\}$. Each value is a proportion over 100 replicates.

| ISE | PF+F | C60+F | C40+F | C30+F | C20+F |
|---|---|---|---|---|---|
| **C60[10]** | 0.0006 | 0.0013 | 0.0184 | 0.0224 | 0.0429 |
| **C60[15]** | 0.0009 | 0.0012 | 0.0173 | 0.0225 | 0.0322 |
| **C60[20]** | 0.0010 | 0.0013 | 0.0155 | 0.0200 | 0.0359 |
| **C60[30]** | 0.0018 | 0.0021 | 0.0093 | 0.0116 | 0.0151 |
| **C60[40]** | 0.0016 | 0.0017 | 0.0060 | 0.0082 | 0.0225 |
| **C60[50]** | 0.0019 | 0.0020 | 0.0063 | 0.0077 | 0.0118 |
| **C60[60]** | 0.0018 | 0.0018 | 0.0046 | 0.0056 | 0.0064 |
| **C60** | 0.0020 | 0.0020 | 0.0036 | 0.0044 | 0.0052 |

Table S4. The mean ISE for data generated under distributions in $\mathfrak{C}60$. This table is computed the same way as Table 1 in the main text but over the distinct generating distributions instead of sequence lengths.

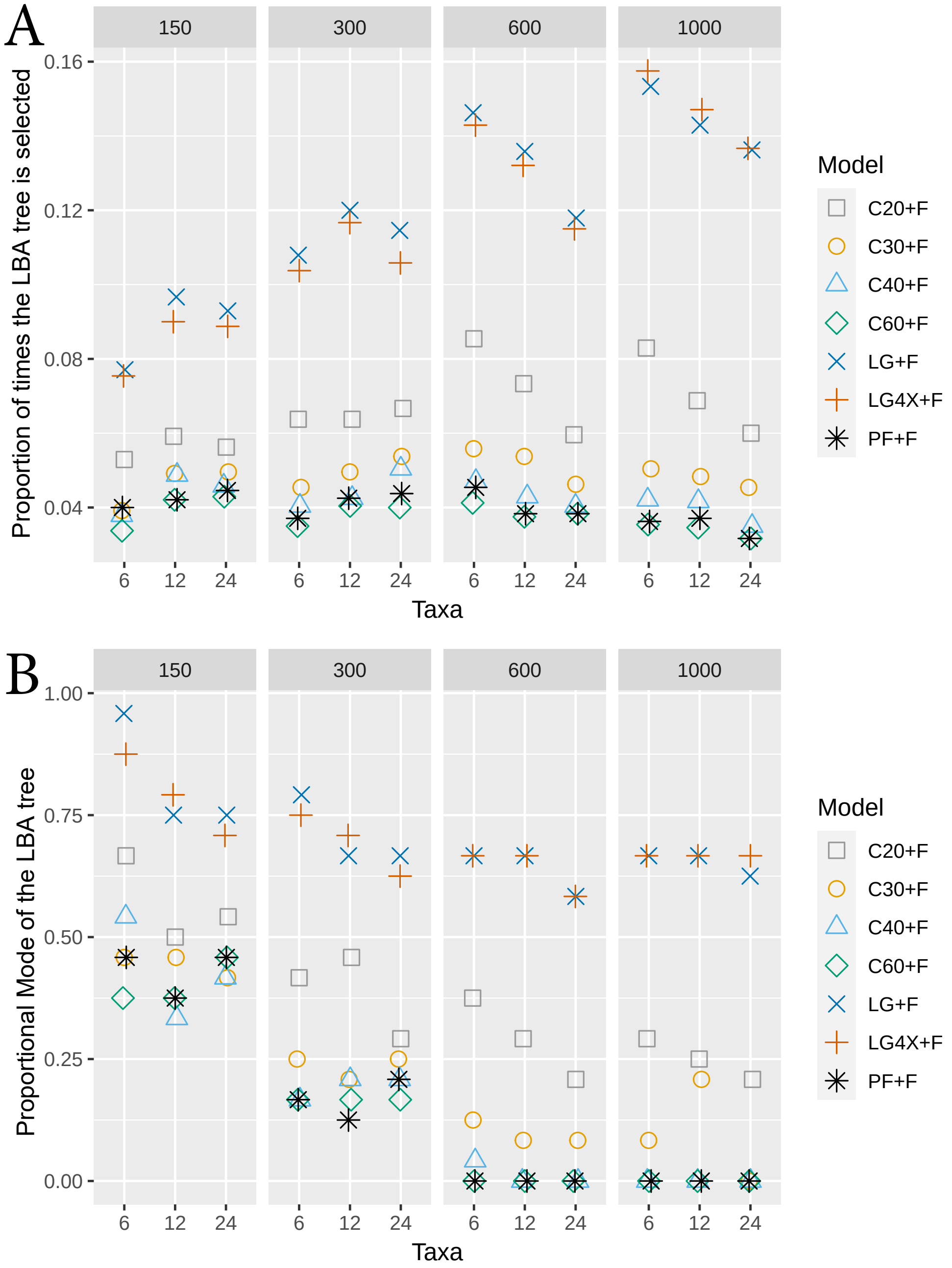


Fig. S2. (A) The plot of the proportion of times the LBA tree is selected for various models fitting data generated under mixing distributions in $\mathfrak{C}60$ fitted to all frequency vectors. Label PF denotes the frequency vectors used to generate the data. The $x$-axis represents the number of taxa on the tree. The plot is divided by the sequence length. (B) A similar plot to that on top but for the proportional model for the LBA tree.

### *Simulations on Bigger Trees*

We performed additional simulations on three trees with more taxa. One of such trees was obtained via a birth-death simulation (40 taxa) and the other two are estimates from empirical data sets (40 and 32 taxa). Details on these trees and the simulations on such are given below.

*Birth-death tree simulation* The tree obtained via a birth-death process, denoted $T_{bd}$, is depicted in Figure S2. This tree was obtained under a Poisson birth-death process on 40 taxa using the R package `castor` (Louca and Doebeli (2017)), with a birth rate of 0.56 and death rate of 0.61. The birth and death rates were chosen so the mean and variance branch length were close to that of the tree $T_I^{20}$, which as explained below is estimated in Wang et al. (2017) from empirical data (data set (I) as referred to in the manuscript). The mean and variance of the branch lengths in $T_I^{20}$ are 0.21 and 0.08, respectively, while for $T_{bd}$ are 0.25 and 0.08.

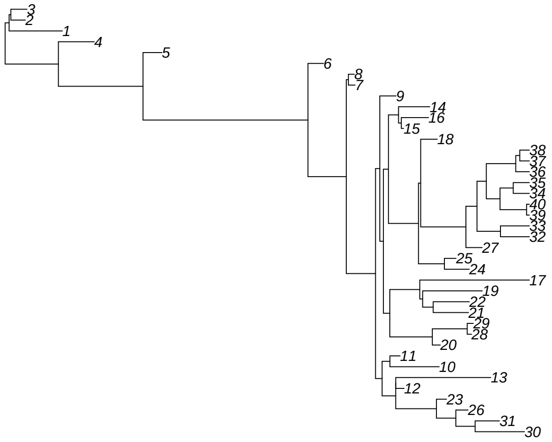


Fig. S3. The tree $T_{bd}$ simulated under a Poisson process on 40 taxa, birth rate 0.56 and death rate 0.61. The Newick notation of such tree is:
((((((((((((30:0.54,31:0.27):0.21,26:0.13):0.21,23:0.11):0.44,(12:0.09,13:1.04):0):0.15,(10:0.54,11:0.11):0.09):0.07,
((((20:0.09,(28:0.05,29:0.07):0.38):0.46,(((21:0.39,22:0.39):0.11,19:0.65):0.03,17:1.2):0.33):0.07,(((24:0.27,25:0.13):0.28,
((27:0.18,((32:0.32,33:0.32):0.26,(((39:0.03,40:0.03):0.29,(34:0.18,35:0.18):0.14):0.15,(36:0.15,(37:0.11,38:0.11):0.04):0.32
):0.1):0.12):0.49,18:0.18):0.02):0.33,((15:0.03,16:0.3):0.03,14:0.34):0.11):0.06):0.04,9:0.18):0.05):0.32,(7:0.07,8:0.06):0.02
):0.42,6:0.17):1.8,5:0.21):0.92,4:0.39):0.58,(1:0.58,(2:0.16,3:0.18):0.02):0.04);

For this tree, we simulated sequences of lengths 100 and 600, POISSON exchangeabilities, $\Gamma(4)$ rates, and frequencies: C60[10], C60[15], and C60[20]. We simulated 50 sequences for each choice of parameters. For all simulations, we did a full tree search on IQ-TREE 2 fitting C60+F, C40+F, C30+F, C20+F, LG4X+F, and POISSON+F. Then we looked at the percentage of correct inferred splits in the tree per simulation. We found a very high percentage of correct inferred splits. Table S5 shows the mean percentage of correct inferred splits in the tree per scenario. Approximately, 79% of the non-trivial splits in the tree are correctly inferred for alignments of length 100 and 94% for alignments of length 600 for all fitted models. Compared to the trees $T_{6m}(l)$ in the manuscript, we see a better percentage of correct inferred splits here than OA values. We believe this is because (1) many of the splits in $T_{bd}$ are not under excessive LBA artifacts; and (2) the more taxa available the better one can infer a split. We see no significant disadvantage in using C60+F versus PF+F when looking the proportion of correct inferred splits. Table S5.5 shows the p-values obtained from a two-sided z-test on the difference between the mean percentage of correct inferred splits with C60 and M, for M$\in$ {C20,C30,C40,PF,LG4X, POISSON}. In congruence with the results in the manuscript, this suggests that when there is no misspecification, over-parameterization is not a problem for complex models. Moreover, this table shows that for very short alignments, C60 does overall significantly better than simpler models (except for C40). Showing also that in some cases there is a significant disadvantage when using simpler models.

Figure S4 shows the box plots for the weight estimates for all classes in C60+F. Here we can see that the true values are, in general, adequately approximated for both alignment lengths. There is, as expected, less variability for the alignments of length of 600. This implies that the observed joint CDF is approaching the true CDF.

| Length | Model | PF+F | C60+F | C40+F | C30+F | C20+F | LG4X+F | POI+F |
|---|---|---|---|---|---|---|---|---|
| 100 | C60[10] | 0.797 | 0.798 | 0.789 | 0.785 | 0.784 | 0.732 | 0.759 |
| | C60[15] | 0.777 | 0.781 | 0.783 | 0.760 | 0.750 | 0.696 | 0.707 |
| | C60[20] | 0.787 | 0.783 | 0.784 | 0.786 | 0.764 | 0.727 | 0.738 |
| 600 | C60[10] | 0.948 | 0.947 | 0.946 | 0.943 | 0.948 | 0.938 | 0.943 |
| | C60[15] | 0.941 | 0.942 | 0.942 | 0.940 | 0.937 | 0.936 | 0.941 |
| | C60[20] | 0.945 | 0.945 | 0.945 | 0.942 | 0.938 | 0.934 | 0.940 |

Table S5. The mean proportion of correct inferred splits estimated for the tree $T_{bd}$ per fitted model under a full tree search. Proportions are over a total of $1850 = 37 \cdot 50$ splits (the tree $T_{bd}$ has 37 non-trivial splits and 50 simulations were performed for each set of frequency classes). POI denotes the POISSON model.

| Length | Model | PF+F | C40+F | C30+F | C20+F | LG4X+F | POI+F |
|---|---|---|---|---|---|---|---|
| 100 | C60[10] | 0.825 | 0.139 | 0.054 | 0.030 | 0.000 | 0.000 |
| | C60[15] | 0.346 | 0.708 | 0.044 | 0.013 | 0.000 | 0.000 |
| | C60[20] | 0.460 | 0.953 | 0.742 | 0.065 | 0.000 | 0.000 |
| 600 | C60[10] | 0.566 | 0.829 | 0.317 | 0.890 | 0.043 | 0.475 |
| | C60[15] | 0.153 | 1.000 | 0.634 | 0.346 | 0.219 | 0.919 |
| | C60[20] | 0.566 | 1.000 | 0.510 | 0.123 | 0.025 | 0.317 |

Table S5.5. The p-values obtained from all the two-sided z-test were we compared the mean percentage of correctly inferred splits between C60 and each alternative model for data simulated under the $T_{bd}$ tree. The null hypothesis is H0: the mean percentage of estimated splits for both models is equal and HA: the means are different. One can determine if C60 performed significantly better than a competing model if (1) the p-value is less than 0.05 and (2) the percentage of correct inferred splits in Table S7 is higher.

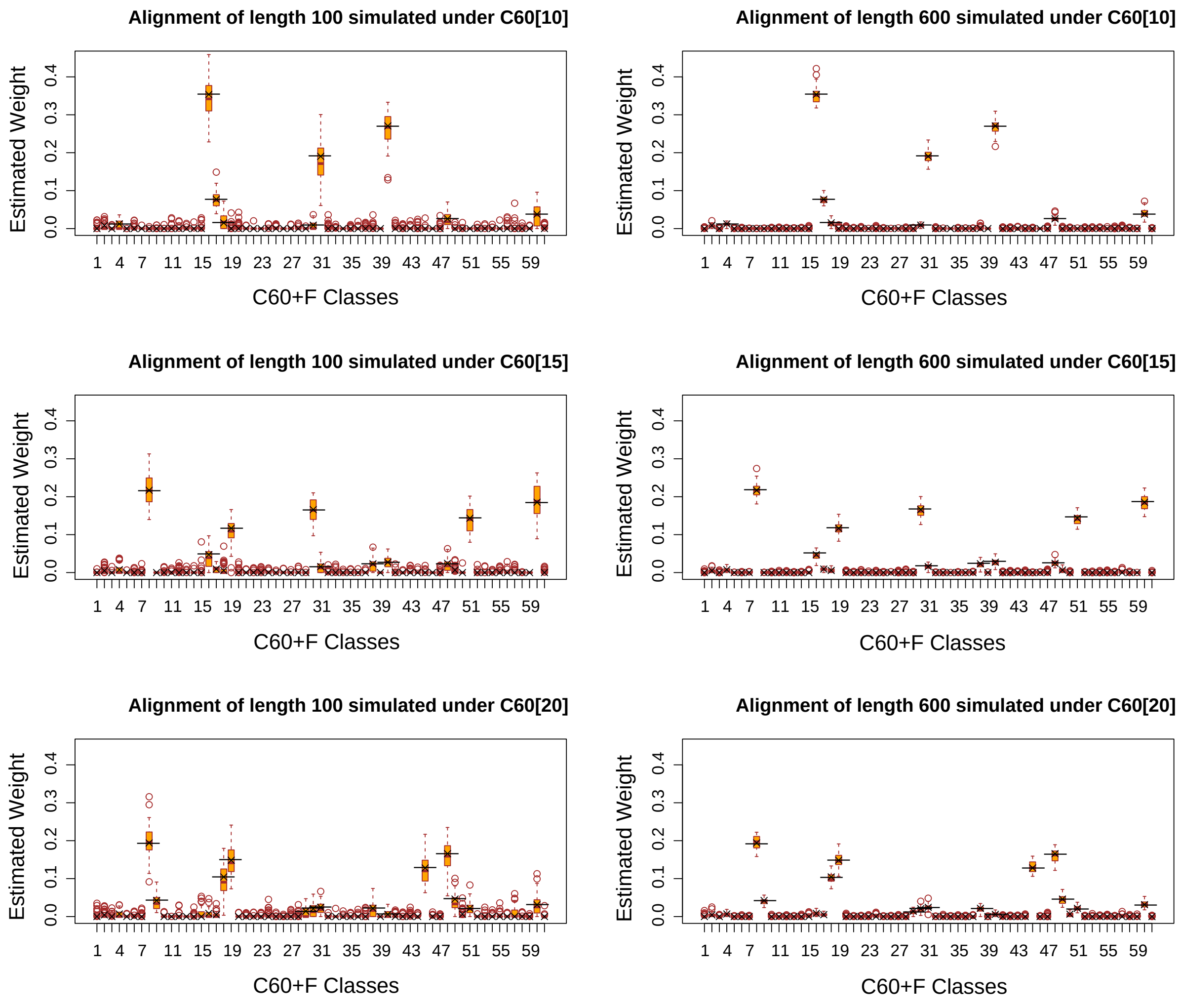


Fig. S4. Estimated weights for the frequency classes in C60+F for all simulations under the tree $T_{bd}$. Each row of plots shows the bar graph for the estimated weights of the C60+F model for data simulated under a fixed model and sequence lengths of 100 and 600. Classes names are in a one-to-one correspondence with the label on the x-axis, except for the F-class which corresponds to label 61. Black lines show the true weights of each class from the true model used to generate the data.

*Empirical trees simulation* The two trees $T_I^{20}$ and $T_{III}^{CAT}$ (as referred to in the manuscript) are the two empirical trees used for simulations. The branch lengths are those estimated in Wang et al. (2017) using the PMSF model. As mentioned in the section *Real Data*, tree $T_I^{20}$ is considered the true topology for data set (I) and $T_{III}^{CAT}$ of data set (III).

Similarly to the tree $T_{bd}$, we simulated sequences of lengths 100 and 600, POISSON exchangeabilities, $\Gamma(4)$ and only model C60[15]. We simulated 50 sequences for each choice of parameters. For all simulations, we did a full tree search on IQ-TREE 2 fitting C60+F, C40+F, C30+F, C20+F, LG4X+F, and POISSON+F. Table S6 shows the percentage of correctly inferred splits for these trees. Similarly as above, Table S6.5 shows the p-values obtained from a two-sided z-test on the difference between the mean percentage of correct inferred splits with C60 and the other models. Similarly as above, we conclude that C60 is not in significantly disadvantage versus the true model. Also, the simpler are in disadvantage versus C60 and C40 performs well.

| Tree | Length | PF+F | C60+F | C40+F | C30+F | C20+F | LG4X+F | POI+F |
|---|---|---|---|---|---|---|---|---|
| $T_I^{20}$ | 600 | 0.953 | 0.951 | 0.952 | 0.944 | 0.938 | 0.888 | 0.901 |
| | 150 | 0.723 | 0.725 | 0.728 | 0.714 | 0.708 | 0.671 | 0.677 |
| $T_{III}^{CAT}$ | 600 | 0.960 | 0.958 | 0.958 | 0.948 | 0.947 | 0.920 | 0.946 |
| | 150 | 0.772 | 0.771 | 0.761 | 0.754 | 0.749 | 0.708 | 0.722 |

Table S6. The mean proportion of correct inferred splits estimated for trees $T_I^{20}$ and $T_{III}^{CAT}$ for sequences generated under the model C60[15]+POISSON. Proportions are over a total of $1850 = 37 \cdot 50$ splits for the tree $T_I^{20}$ (such tree has 37 non-trivial splits) and $1450 = 29 \cdot 50$ splits for the tree $T_{III}^{CAT}$ (such tree has 29 non-trivial splits). POI denotes the POISSON model.

| Tree | Length | PF+F | C40+F | C30+F | C20+F | LG4X+F | POI+F |
|---|---|---|---|---|---|---|---|
| $T_I^{20}$ | 600 | 0.194 | 0.823 | 0.255 | 0.038 | 0.000 | 0.000 |
| | 150 | 0.448 | 0.674 | 0.124 | 0.065 | 0.000 | 0.000 |
| $T_{III}^{CAT}$ | 600 | 0.077 | 1.000 | 0.017 | 0.007 | 0.000 | 0.035 |
| | 150 | 0.867 | 0.144 | 0.038 | 0.011 | 0.000 | 0.000 |

Table S6.5. The p-values obtained from a two-sided z-test were we compared the mean percentage of correctly inferred splits between C60 and the each alternative model, for data simulated under trees $T_I^{20}$ and $T_{III}^{CAT}$ and the model C60[15]+POISSON. The null hypothesis is H0: the mean percentage of estimated splits for both models is equal and HA: the means are different. One can determine if C60 performed significantly better than a competing model if (1) the p-value is less than 0.05 and (2) the percentage of correct inferred splits in Table S8 is higher.

To focus on the effects of LBA artifacts known in the data sets of these trees, we also simulated alignments of lengths 300, 600, and 1000 using all sets of frequency classes in $\mathfrak{C}60$. In all cases, POISSON exchangeabilities were used together with $\Gamma(4)$ rates with $\alpha = 0.5$. For each scenario, 50 simulations were created. Then we compared the fit of the true and alternative topologies (the LBA topology) for each of these trees ($T_I^{LG}$ and $T_{III}^{LG}$ for $T_I^{20}$ and $T_{III}^{CAT}$, respectively). Table S7 shows the number of times the true topology was preferred over the alternative. We again see a better performance of complex models. Note that these percentages are in the vicinity of the percentage of correct inferred splits. This reinforces the fact that LBA is the force biasing inference.

We also computed the mean ISE for data estimated under the true trees. These results are shown in Table S8. We see that complex models have better ISE values. In all cases, ISE decreases as alignment length increases. All these results are aligned with the findings in the manuscript and the simulations on the $T_{bd}$ tree.

| | 300 | | 600 | | 1000 | |
|---|---|---|---|---|---|---|
| | $T_I^{20}$ | $T_{III}^{CAT}$ | $T_I^{20}$ | $T_{III}^{CAT}$ | $T_I^{20}$ | $T_{III}^{CAT}$ |
| PF+F | 0.94 | 0.70 | 0.99 | 0.87 | 0.99 | 0.93 |
| C60+F | 0.94 | 0.71 | 0.99 | 0.87 | 0.99 | 0.92 |
| C40+F | 0.93 | 0.67 | 0.99 | 0.86 | 0.99 | 0.92 |
| C30+F | 0.92 | 0.68 | 0.99 | 0.84 | 0.99 | 0.90 |
| C20+F | 0.91 | 0.63 | 0.99 | 0.78 | 0.99 | 0.84 |
| LG4X | 0.75 | 0.46 | 0.86 | 0.52 | 0.90 | 0.53 |
| POI+F | 0.79 | 0.52 | 0.91 | 0.61 | 0.92 | 0.63 |

Table S7. The proportion of times the (true) trees $T_I^{20}$ and $T_{III}^{CAT}$ were preferred over their LBA tree counterparts for the data generated under models $\mathfrak{C}60$ and POISSON exchangeabilities.

| | | ISE | |
|---|---|---|---|
| Length | Fitted | $T_I^{20}$ | $T_{III}^{CAT}$ |
| 300 | PF+F | 0.00089 | 0.00122 |
| | C60+F | 0.00092 | 0.00129 |
| | C40+F | 0.00954 | 0.00976 |
| | C30+F | 0.01142 | 0.01169 |
| | C20+F | 0.01317 | 0.01340 |
| 600 | PF+F | 0.00045 | 0.00060 |
| | C60+F | 0.00046 | 0.00063 |
| | C40+F | 0.00913 | 0.00919 |
| | C30+F | 0.01102 | 0.01118 |
| | C20+F | 0.01271 | 0.01283 |
| 1000 | PF+F | 0.00027 | 0.00038 |
| | C60+F | 0.00028 | 0.00040 |
| | C40+F | 0.00895 | 0.00903 |
| | C30+F | 0.01084 | 0.01104 |
| | C20+F | 0.01254 | 0.01259 |

Table S8. The mean ISE obtained from fitting the (true) trees $T_I^{20}$ and $T_{III}^{CAT}$ for data generated under models $\mathfrak{C}60$ and POISSON exchangeabilities.

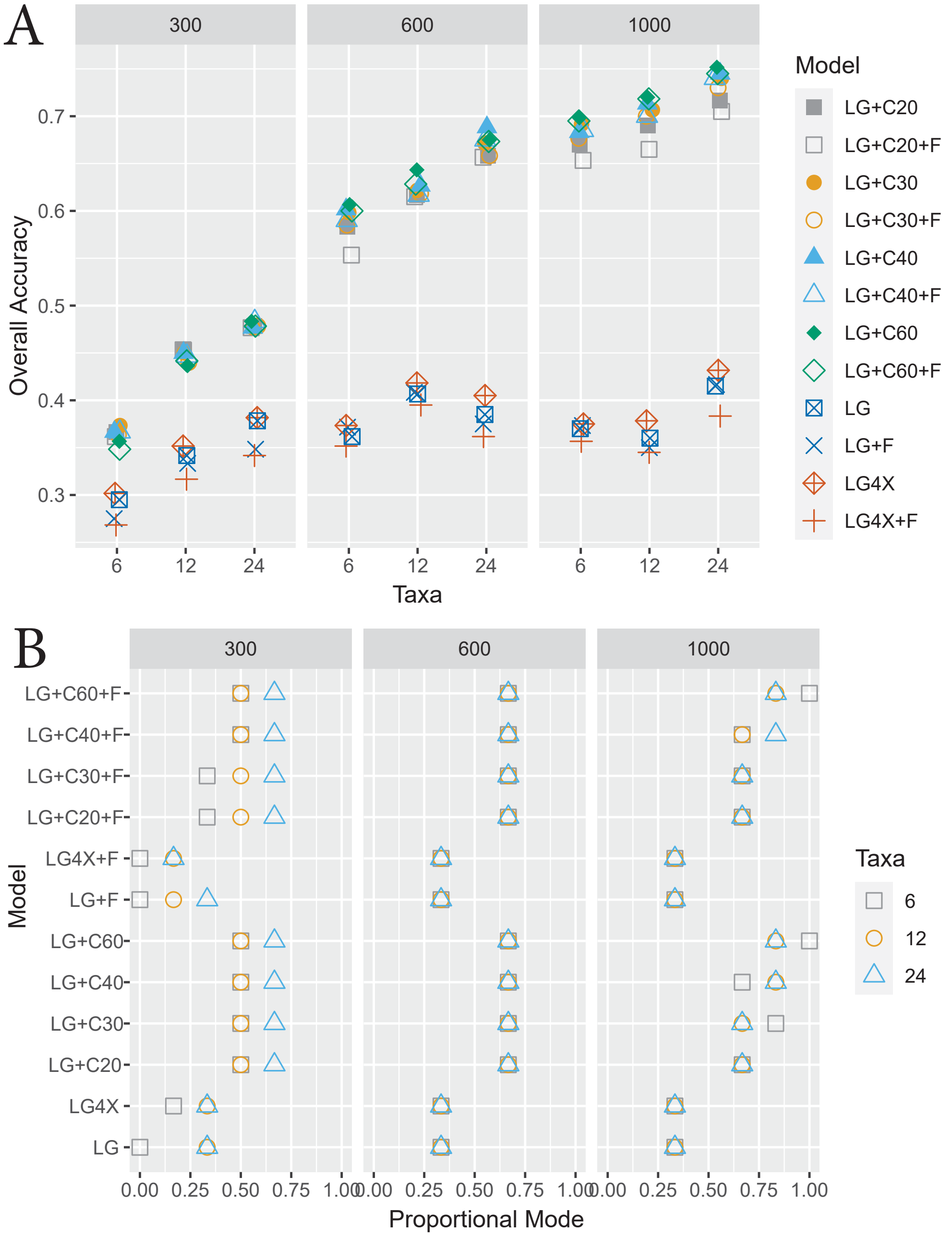


Fig. S5. (A) The plot of the OA values per model of data generated under the UDM distributions and LG exchangeabilities. Different classes are fitted but in all cases we fit LG exchangeabilities. The $x$-axis represents number of taxa on the tree. The plot is divided by the sequence length. (B) A similar plot to that on top but for PM.

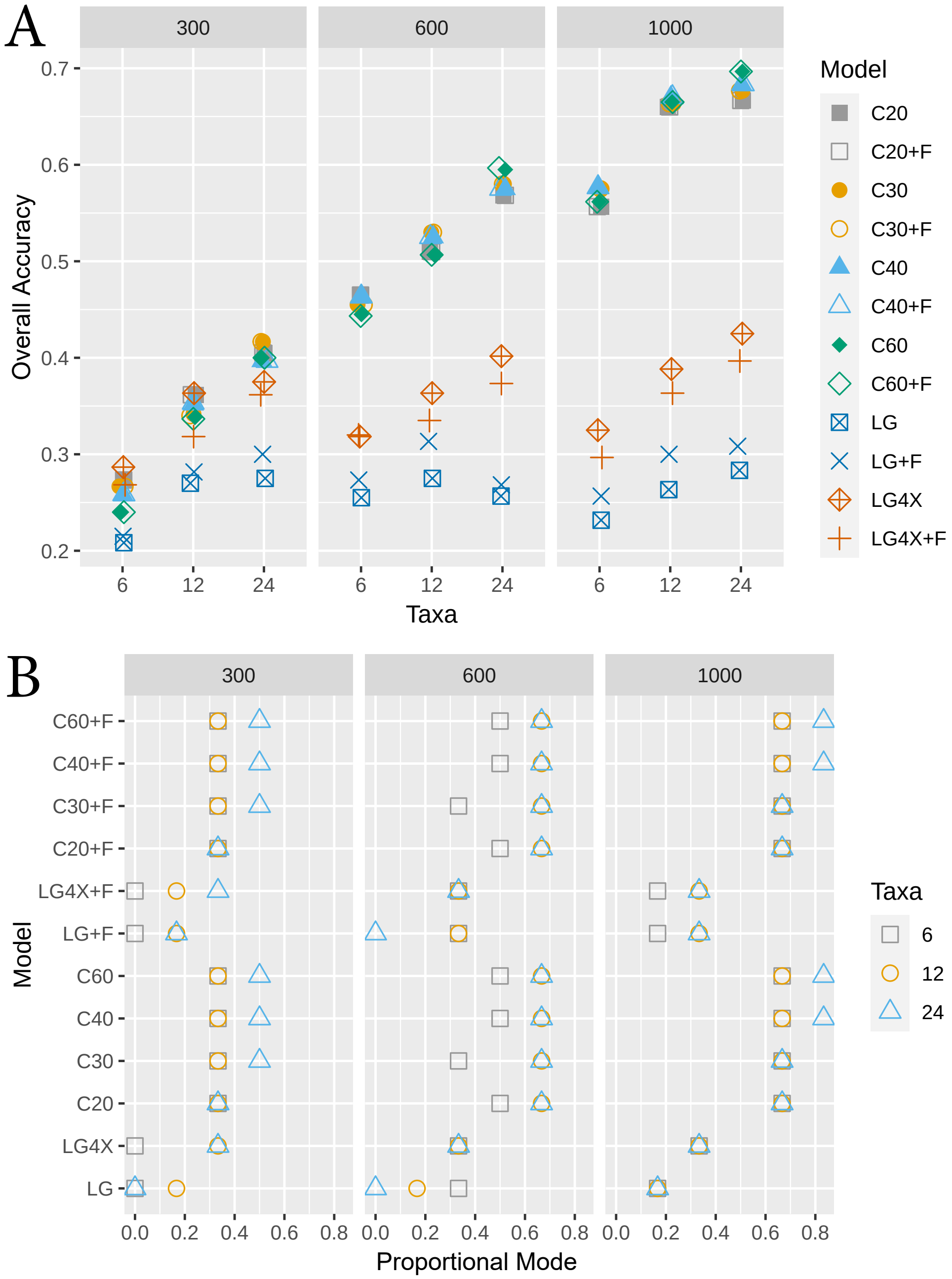


Fig. S6. (A) The plot of the OA values per model of data generated under the UDM distributions and LG exchangeabilities. Different classes are fitted but in all cases we fit POISSON exchangeabilities. The $x$-axis represents number of taxa on the tree. The plot is divided by the sequence length. (B) A similar plot to that on top but for PM.

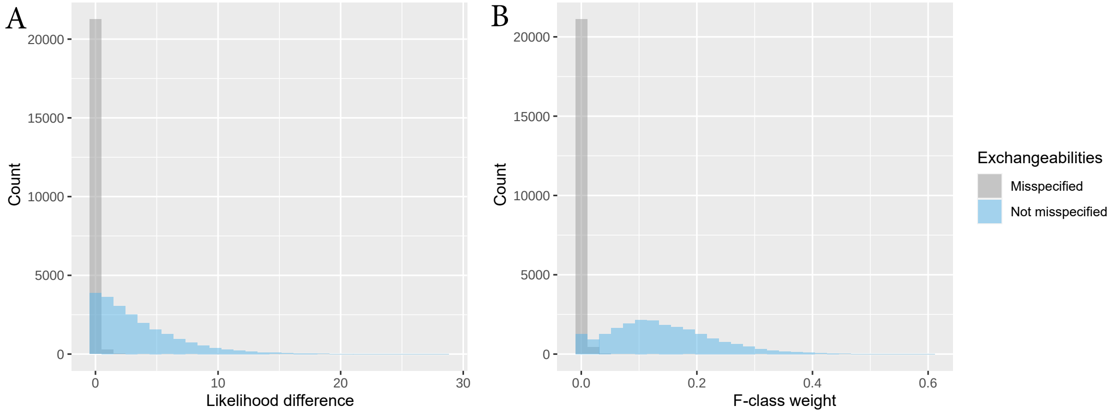


Fig. S7. (A) Histograms showing the difference between likelihoods values of models fitted with and without the F-class. No misspecification of the exchangeabilities is depicted in blue and in gray when there is. (B) Histograms showing the inferred F-class weight when there is no misspecification of the exchangeabilities (blue) and when there is (gray). The histograms consist of data generated under all trees, number of taxa, both UDM mixing distributions, and LG exchangeabilities. Misspecification of exchangeabilities refers to fitting using POISSON matrix instead of the LG matrix.

#### *F-class bias investigation via the Shannon entropy*

As mentioned in the text, we explored the bias observed in weight of the F-class when there is misspecification of the exchangeabilities. This bias is investigated in relation to the uniformity of frequencies at sites as measured by Shannon entropy.

The Shannon entropy, as defined in our context

$$H(\boldsymbol{\pi}) = -\sum_{j=1}^{20} \pi_j \ln(\pi_j),$$

is a common measure of the degree of uniformity of the amino acid frequencies at sites. We note that when there is misspecification of the exchangeabilities, there is a bias towards frequency classes with high entropy. For data simulated using the POISSON matrix and fitted using the LG matrix, more than 80% of the 27000 data sets where we fitted a model with the F-class, this was the class with the highest entropy. When the F-class had the highest entropy, it was assigned, on average, more than half the total weight (average weight = 0.59). Moreover, when fitting without the F-class, we noted that, in general, the class with the highest entropy is assigned a really large weight. For example, for all 5400 simulations (9 trees, 3 sequence lengths, 2 UDM distributions, and 100 repetitions per condition) when fitting the classes in either C20, C30, or C60, the class with the highest entropy had the largest weight, and on average, that weight was 4.71 times more than the weight assigned to that class when there is no misspecification. When fitting the classes in C40, the class with the second-highest entropy is the one with the largest weight, and it was also, on average, 3.99 times more than its weight when there is no misspecification. Therefore we observe that this bias may not directly related to entropy but may instead be some other factor that is correlated with entropy.

Although, for data generated using the LG matrix but fitted the POISSON matrix, we found a shift in the correlation of entropy and class weight for all models. The average correlation between entropy and class weight for C60, C40, and C30 is 0.21 when there is no misspecification and -0.56 when there is. For C20 the shift is in a different direction, i.e the correlation between entropy and class weight is -0.12 when there is no misspecification and 0.11 when there is. Nonetheless, for this model and when there is misspecification, the class with the highest entropy is the one with the second lowest weight (average weight = 0.008). This same class has the highest weight when there is no misspecification (average weight = 0.142). This also suggests that entropy is somehow related to or affected by, model misspecification.

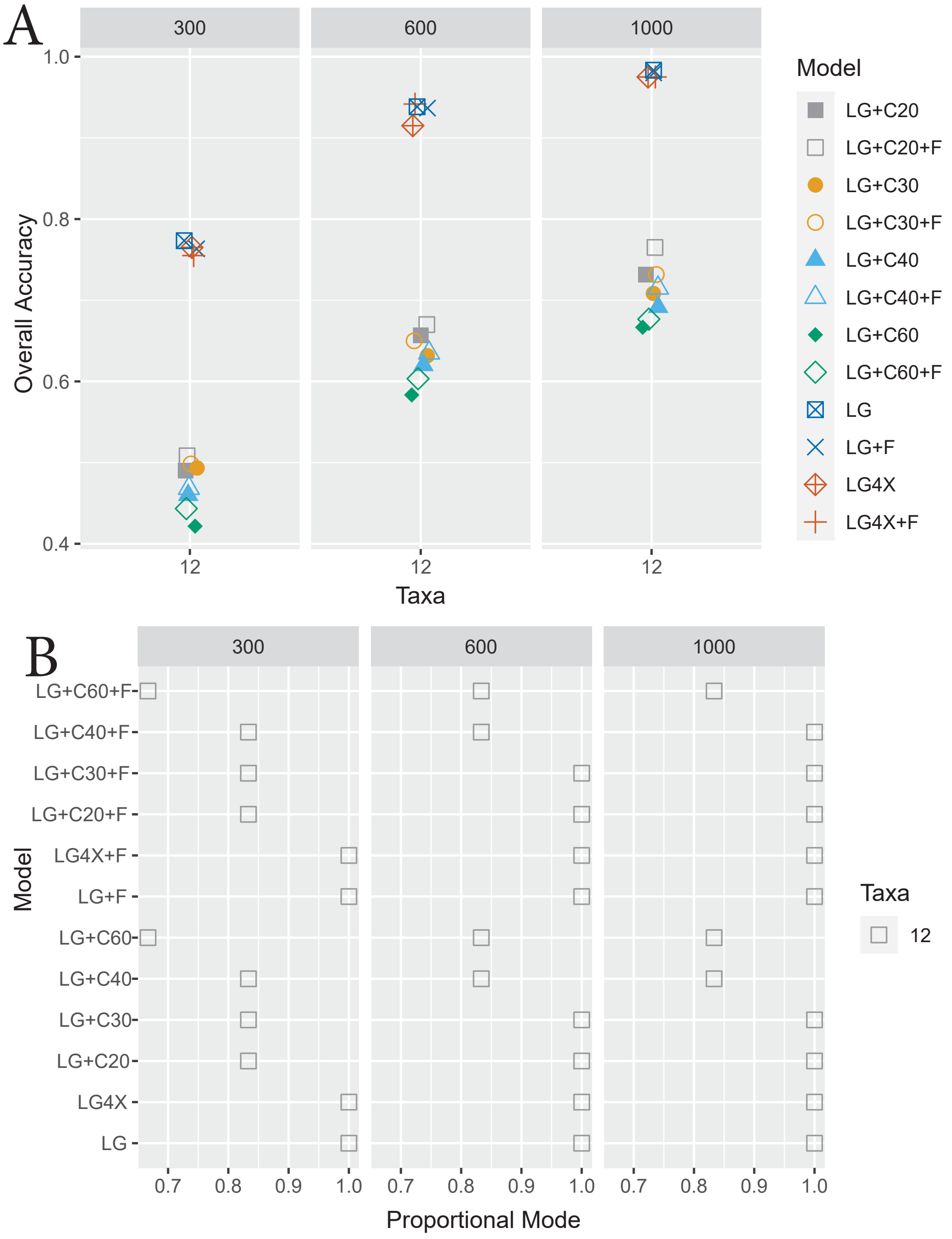


Fig. S8. (A) The plot of the OA values per model of data generated under the UDM distributions and LG exchangeabilities. The trees used to generate the data have long branches apart. They differ from the trees in Figure 1 in having the $e$ and $f$ taxa together rather than separated. Specifically, the trees are: (((f1:.1,f2:.1):1,(e1:.1,e2:.1):1):$l$,(((c1:.1,c2:.1):0.1,(d1:.1,d2:.1):0.1):0.1, ((a1:.1,a2:.1):0.1,(b1:.1,b2:.1):0.1):0.1):$l$)r; with $l \in \{0.05, 0.02, 0.005\}$. Different classes are fitted but in all cases we fit LG exchangeabilities. The plot is divided by the sequence length. (B) A similar plot to that on top but for PM.

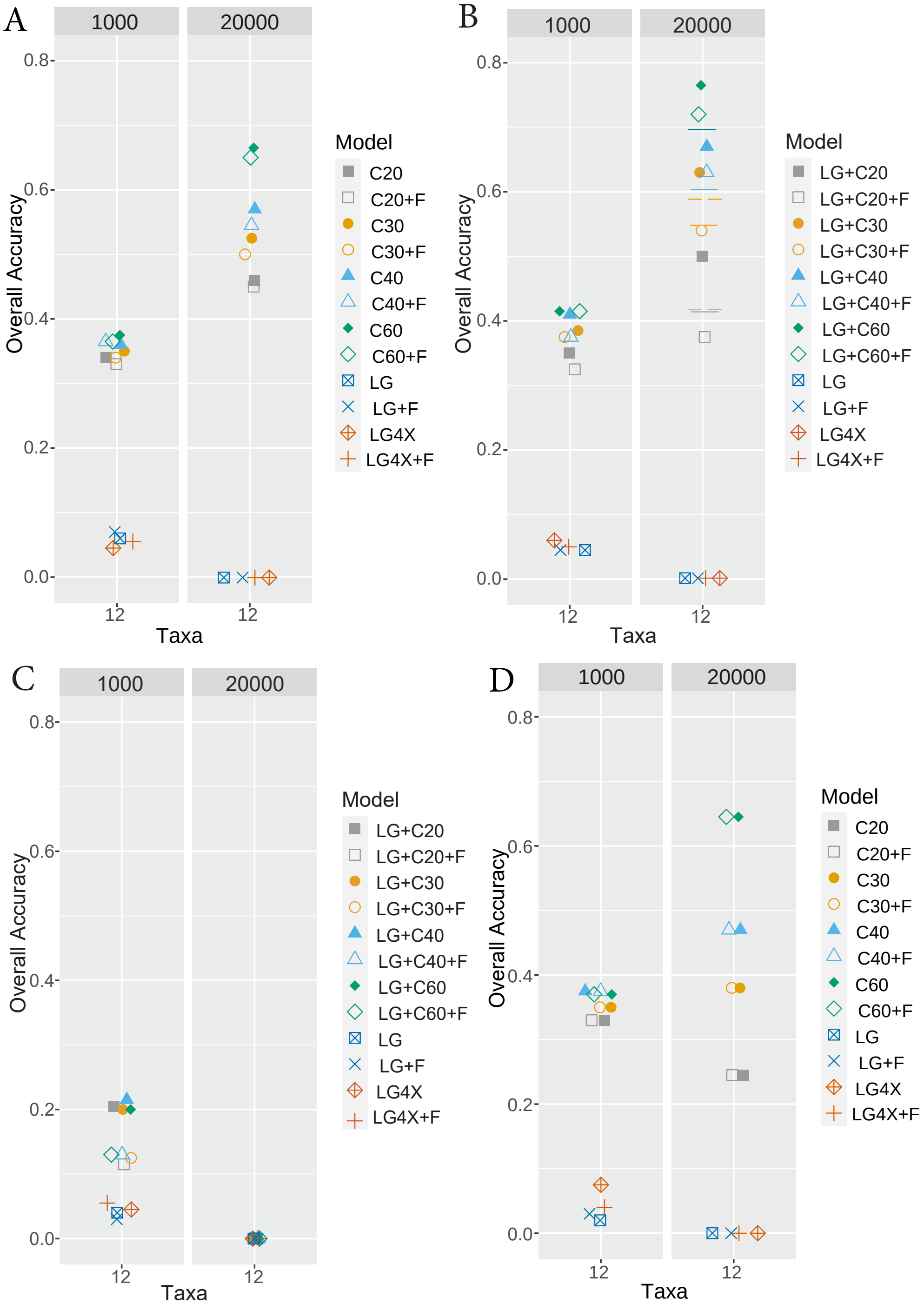


Fig. S9. The plot of the overall accuracy, OA values per model of data generated under the UDM distributions, tree $T_{12}(0.005)$, alignment lengths of 1000 and 20,000, and different exchangeabilities. (A) Shows data generated under POISSON exchangeabilities and fitting using the same. (B) Shows data generated under LG exchangeabilities and fitting the same. (C) Shows data generated under POISSON exchangeabilities and fitting LG. (D) Shows data generated under LG exchangeabilities and fitting POISSON. Each dot in a plot represents 200 simulations (2 UDM models and 100 repetitions of each).

#### *Reproducibility*

The exact version of IQ-tree2 used was `IQ-TREE multicore version 2.1.0 COVID-edition`. This version contains `Alisim`. An example of an `Alisim` routine used in this work, particularly, one showing the simulation of an alignment of length 1000 under the model LG+UDM4096NONE+$\Gamma(0.5)$ and the tree on 12 taxa $T_{12}(0.05)$ described in the manuscript, is the following:

```
iqtree --alisim UDM_LG_1000_wt2_rep98_T12_m005 -mdef
udm_hogenom_4096_none_iqtree.nex  -m LG+UDM4096NONE+G4{0.5}
-t T12_m005.txt --seqtype AA   --length 1000 -af fasta
--skip-checking-memory
```

An example of the `iqtree` routine used in this work is shown below, particularly one fitting the model POISSON+C40+$\Gamma(0.5)$ to all 35 topologies shown in table S1, for the alignment obtained from `Alisim` described above, is the following:

```
iqtree -s UDM_LG_1000_wt2_rep98_T12_m005_0.fa -me .00001
-m POISSON+C40+G -z 12TaxafileTopo35.txt -nt 2
-pre C40noF_P_LG_1000_wt2_98_T12_m002 -a 0.5 -te T12_m002.txt -mwopt
```

While `Alisim` is a very powerful simulator, for this case study, we focus on simple sequence simulation strictly on profile mixture models. Therefore our simulations do not contain insertions nor deletions. The simulations process is exactly as expected from the description of profile mixture models in Section *Mixture models and over-parameterization with large samples* in the manuscript.
